# Extreme Mitogenomic Variation Without Cryptic Speciation in Chaetognaths

**DOI:** 10.1101/025957

**Authors:** Ferdinand Marlétaz, Yannick Le Parco, Shenglin Liu, Katja T.C.A. Peijnenburg

## Abstract

The extent of within-species genetic variation across the diversity of animal life is a fundamental but largely unexplored problem in ecology and evolution. The neutral theory of molecular evolution predicts that genetic variation scales positively with population size. However, the genetic diversity of mitochondrial DNA, a prominent marker used in DNA barcoding studies, shows very little variation across animal species. Here, we report an unprecedented case of extreme mitochondrial variation within natural populations of two species of chaetognaths (arrow worms). We determined that this diversity is composed of deep intraspecific mitochondrial lineages within single populations that could be as divergent as human and newt. This mitochondrial diversity is the highest ever reported in animals without evidence of cryptic speciation or allopatric divergence as supported by nuclear evidence. We sequenced 54 complete mitogenomes revealing gene order rearrangements between these intraspecific lineages. Such structural differences have never previously been reported within single species. We confirm that this divergence was not driven by positive selection, and conversely show that these lineages evolved under purifying selection, consistently with neutral expectations. Our findings question the generally accepted narrow range of genetic variation in animal mitochondria and argue for a reappraisal of DNA barcoding techniques. Furthermore, extreme levels of mitogenomic variation in chaetognaths challenge classical views regarding mitochondrial evolution and cytonuclear co-evolution.

## 1. Introduction

Genetic diversity estimates provide information about a species’ population history, dynamics, and adaptive potential (*e.g.* [1-3]). Mitochondrial genes have been widely used to characterize natural populations at the molecular level, mostly because of technical ease-of-use considerations such as clonality and high mutation rate [4]. In particular, the DNA barcoding approach has propelled the use of mitochondrial gene fragments, such as Cytochrome Oxidase 1 (Cox1). The combination of strong interspecific divergence and generally low levels of intraspecific variation, also known as the barcoding ‘gap’, makes such markers an essential tool for species identification [5]. Many observations of high levels of mitochondrial divergence have consequently been interpreted as evidence of cryptic speciation, the distinct genetic entities being often, but not always, associated with previously overlooked geographic, morphological, or reproductive boundaries [1].

The observation that the range of genetic variation does not scale with population size as predicted by the neutral theory of molecular evolution was referred to by Lewontin as the ‘paradox of variation’ [6,7]. This paradox is particularly manifest for animal mitochondrial diversity levels, which show surprisingly little variation across species [8]. This has been interpreted as the consequence of pervasive positive selection [8], genetic hitch-hiking [9], or as the result of an inverse correlation between population size and mutation rate [10]. Similarly, mitochondrial transplantation experiments have demonstrated that the cyto-nuclear interactions in respiratory complexes are an important factor limiting diversity in mitochondrial genes [11,12]. Nevertheless, species with large population size are expected to simultaneously show high genetic diversity and strong purifying selection, a combination that has never been observed in nature for mitochondria [13]. However, most of the data gathered to assess mitochondrial evolution in animals has come from a subset of well-studied animal groups, such as mammals and insects [8].

The diversity of oceanic animals and their genetics remain very poorly characterized [3,5]. Chaetognaths are an enigmatic marine phylum, which possesses a unique combination of morphological, developmental and genomic characters. They occupy a remarkable phylogenetic position among bilaterian animals as an early protostome lineage [14-17]. They also represent a major planktonic group and play important roles in marine food webs as the primary predators of copepods [18]. The first papers examining the genetics of planktonic chaetognaths reported high levels of diversity and significant population structuring, but also revealed unusual patterns of mitochondrial evolution [19,20]. In this study, we focus on single populations of a benthic (*Spadella cephaloptera*) and two planktonic (*Sagitta elegans* and *Sagitta setosa*) chaetognaths. We intensely sampled these populations to characterize mitochondrial and nuclear diversity and we sequenced whole mitochondrial genome in a large number of in individuals. We show that extreme levels of mitogenomic variation can exist within species without apparent evolutionary consequences.

## 2. Results

### (a) Extreme mitochondrial diversity in two chaetognath species

We uncovered extreme levels of mitochondrial diversity in distinct natural populations of the species *Spadella cephaloptera* and *Sagitta elegans*, two members of Chaetognatha (arrow worms). We conducted extensive genotyping for several mitochondrial markers (Cox1, Cox2 and 16S) of individual chaetognaths sampled at single geographic localities in Sormiou (France) and Gullmar fjord (Sweden) (Table 1, Figure S1). We found that respective nucleotide diversities (π) for Cox1 of *Spadella cephaloptera* (*N*=25) and *Sagitta elegans* (*N*=107) were 0.14 and 0.18. In both species, phylogenetic analysis of individual mitochondrial genes revealed that this diversity is distributed across 8 to 11 highly divergent lineages, with more than 10% uncorrected sequence divergence (Figure 1 and Figure S2). However, we did not detect such high levels of mitochondrial nucleotide diversity in a third species, *Sagitta setosa* (N=54), which is considered a closely related species to *Sagitta elegans*, and was sampled at the same location in Gullmar fjord (Table 1).

**Table 1.**
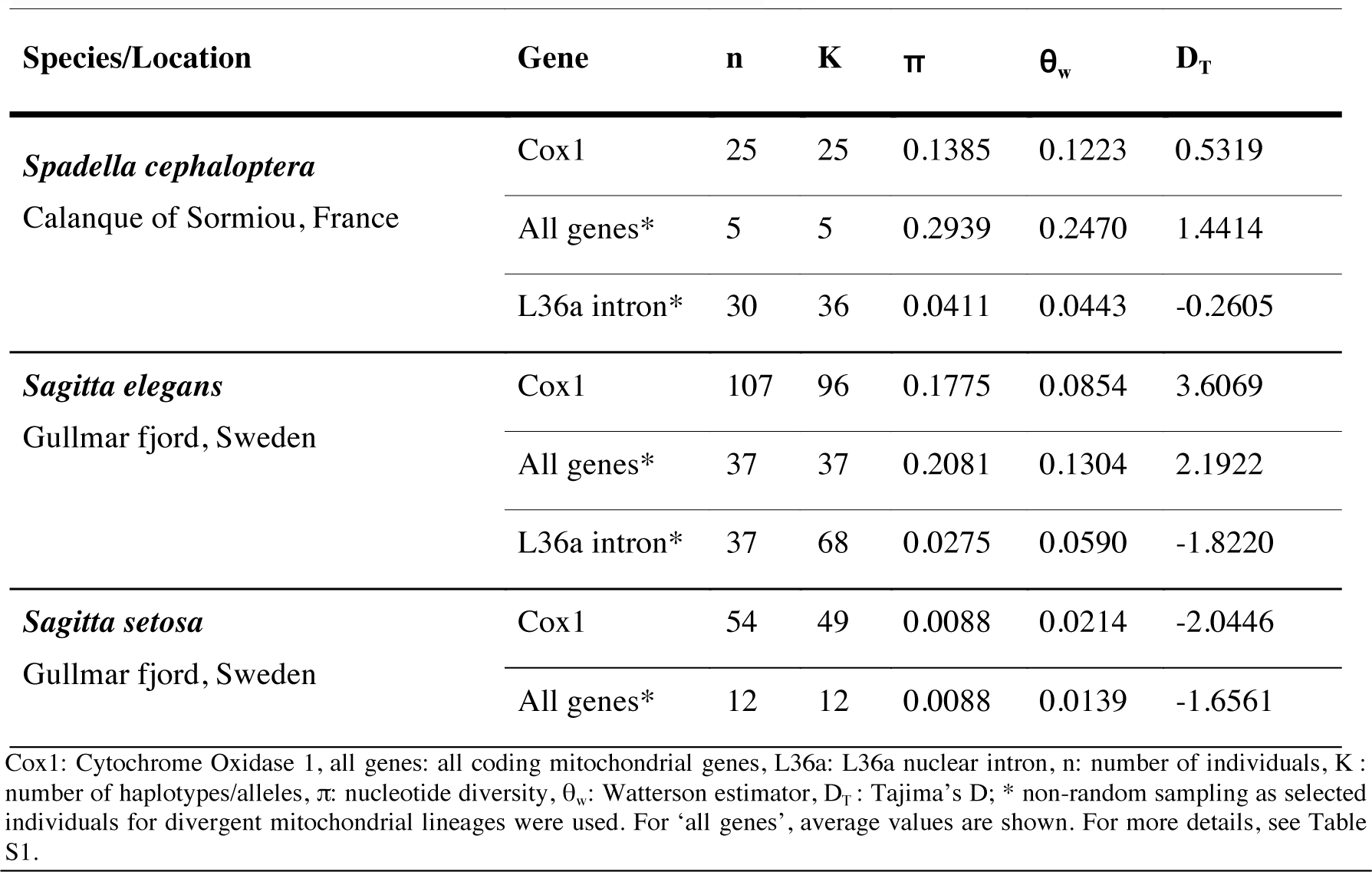
Genetic diversity estimates for selected loci in single populations of the chaetognaths *Spadella cephaloptera, Sagitta elegans*, and *S. setosa*.

| Species/Location | Gene | n | K | $\pi$ | $\theta_w$ | $D_T$ |
| --- | --- | --- | --- | --- | --- | --- |
| *Spadella cephaloptera* | Cox1 | 25 | 25 | 0.1385 | 0.1223 | 0.5319 |
| Calanque of Sormiou, France | All genes* | 5 | 5 | 0.2939 | 0.2470 | 1.4414 |
|  | L36a intron* | 30 | 36 | 0.0411 | 0.0443 | -0.2605 |
| *Sagitta elegans* | Cox1 | 107 | 96 | 0.1775 | 0.0854 | 3.6069 |
| Gullmar fjord, Sweden | All genes* | 37 | 37 | 0.2081 | 0.1304 | 2.1922 |
|  | L36a intron* | 37 | 68 | 0.0275 | 0.0590 | -1.8220 |
| *Sagitta setosa* | Cox1 | 54 | 49 | 0.0088 | 0.0214 | -2.0446 |
| Gullmar fjord, Sweden | All genes* | 12 | 12 | 0.0088 | 0.0139 | -1.6561 |

**Figure 1.**
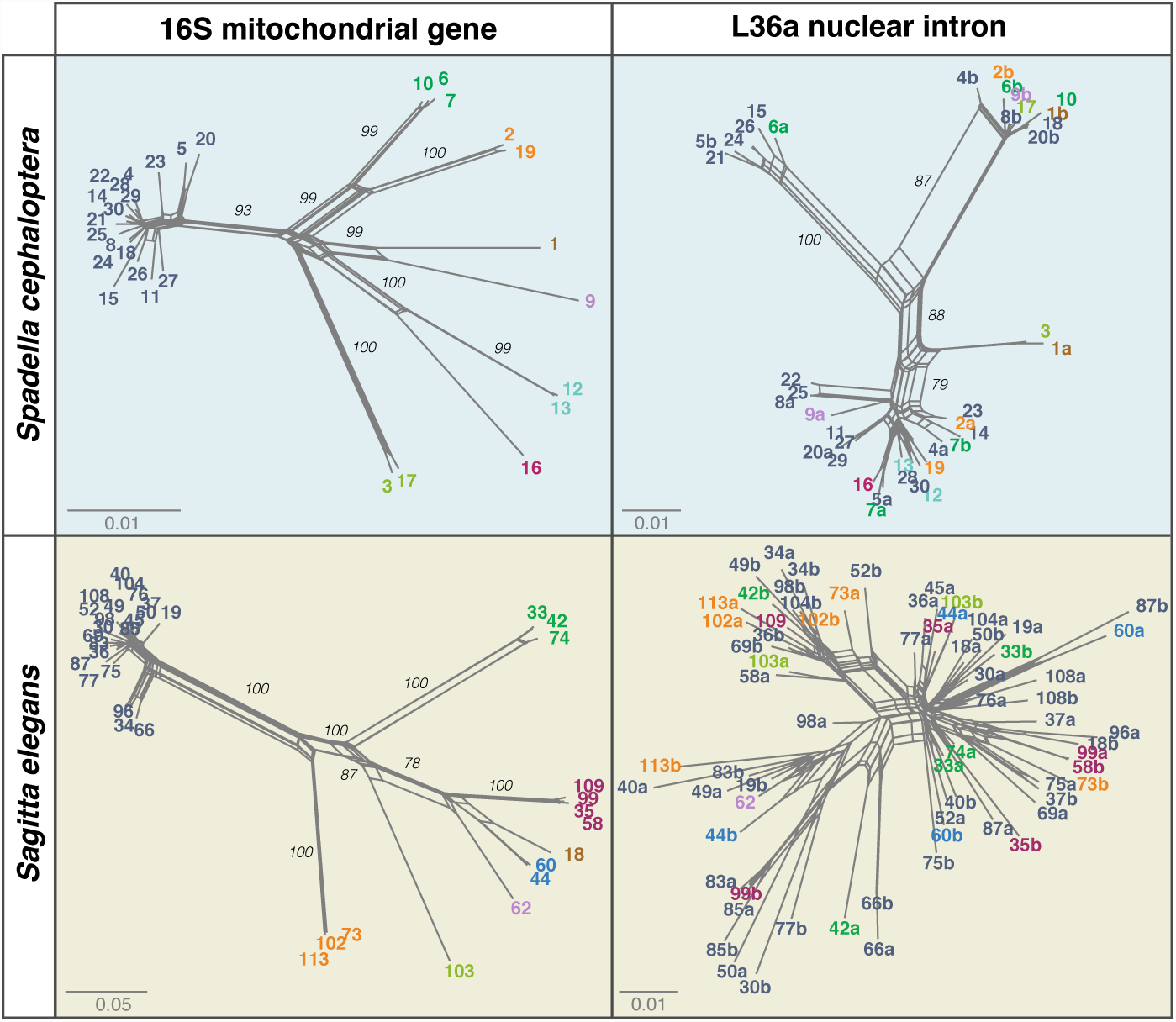
Incongruence of mitochondrial and nuclear lineages in single populations of chaetognaths *Spadella cephaloptera* and *Sagitta elegans*. Neighbor-net phylogenetic networks of mitochondrial 16S (left) and nuclear L36a intron (right) sequences reconstructed using K2P genetic distances. Numbering of individuals is the same for the two markers in each species and labels are colored according to mitochondrial lineage assignment. For nuclear sequences, a and b denote two different alleles recovered by cloning from the same individual. Scale represents expected nucleotide changes per site.

We sequenced mitochondrial gene fragments from >180 individual chaetognaths, but never observed consistent double Sanger chromatogram peaks, which would be indicative of multiple mitochondrial copies [21] or nuclear pseudogenes [22]. We further characterized these lineages by sequencing complete mitochondrial genomes for representative individuals in all three species. We amplified by PCR 54 whole mitochondrial genomes and all amplicons included the 13 genes typical of chaetognath mitochondrial genomes (see Methods). We also did not find any stop codons or frameshift mutations despite global nucleotide diversities of 0.29 and 0.21 in *Spadella cephaloptera* and *Sagitta elegans*, respectively (Table 1). These multiple lines of evidence reject heteroplasmy and pseudogenization hypotheses as an explanation for the extreme levels of diversity observed here.

### (b) Deep mitochondrial lineages do not represent cryptic species

We tested for the presence of cryptic species by comparing the patterns of mitochondrial divergence with those inferred from three nuclear markers: the variable ribosomal protein L36a intron (Figure 1) and the more conserved 18S and 28S rRNA genes frequently used for species delimitation (Table S1). If deep mitochondrial lineages represent distinct, reproductively isolated, species, then we expect to see congruent divergence patterns across independently evolving loci. Conversely, we observed complete incongruence between the topologies obtained from mitochondrial and nuclear markers for both species (Figure 1). In *Spadella cephaloptera*, nuclear intron sequences arrange in several clades, which each include individuals and alleles belonging to distinct mitochondrial lineages (Figure 1). In *Sagitta elegans*, we found significant levels of recombination in the nuclear intron dataset, as would be expected for an interbreeding population of individuals (phi-test, p-value=0). Similarly, 18S and 28S loci show very little variation and no phylogenetic pattern (0 variable sites in *S. elegans* and 2.6% in *S. cephaloptera*, Table S1 and Dataset S1). While these markers can show little variation between species in other groups, they exhibit significant interspecific divergence in chaetognaths and have thus have been extensively used to resolve intraphyletic relationships [23-25]. In sum, we considered three nuclear loci with varying evolutionary rates and none showed patterns of divergence congruent with the observed mitochondrial lineages in either of two species. Hence, we conclude that highly divergent mitochondrial lineages are present within interbreeding populations of *Spadella cephaloptera* and *Sagitta elegans*.

### (c) Class-level divergence between intraspecific lineages

Mitochondrial gene trees for each species and their concatenation from 54 individual mitogenomes show congruent topologies as expected for a haploid, non-recombining genome (Dataset S2). We thus examined phylogenetic relationships and divergence using the complete set of mitochondrial protein-coding sequences along with existing mitogenomic data from other chaetognath species (Figure 2*a*) [26-28]. To put mitochondrial divergence levels into perspective, we compared the phylogenetic distance between chaetognath mitochondrial lineages with those inferred between the main vertebrate groups (Figure 2*b*). We found that distances between mitochondrial lineages of *Spadella cephaloptera* and *Sagitta elegans* compare with those recovered within amniote and mammalian lineages, which diverged 312 and 170 Myr ago, respectively [29]. We assessed the maximum divergence dates compatible with these phylogenetic distances by performing molecular dating analyses. Using calibrations based on geological events (see Methods), we recovered an oldest putative origin of 80 and 180 Myr ago for the divergent intraspecific mitochondrial lineages in *Spadella cephaloptera* and *Sagitta elegans*, respectively (Figure S4). These analyses predict a faster evolutionary rate in chaetognaths than in vertebrates (0.0059±0.0021 versus 0.0024±0.0011 substitutions per site per Myr), but are also compatible with a more recent origin of chaetognath lineages associated with an even faster mutation rate.

**Figure 2.**
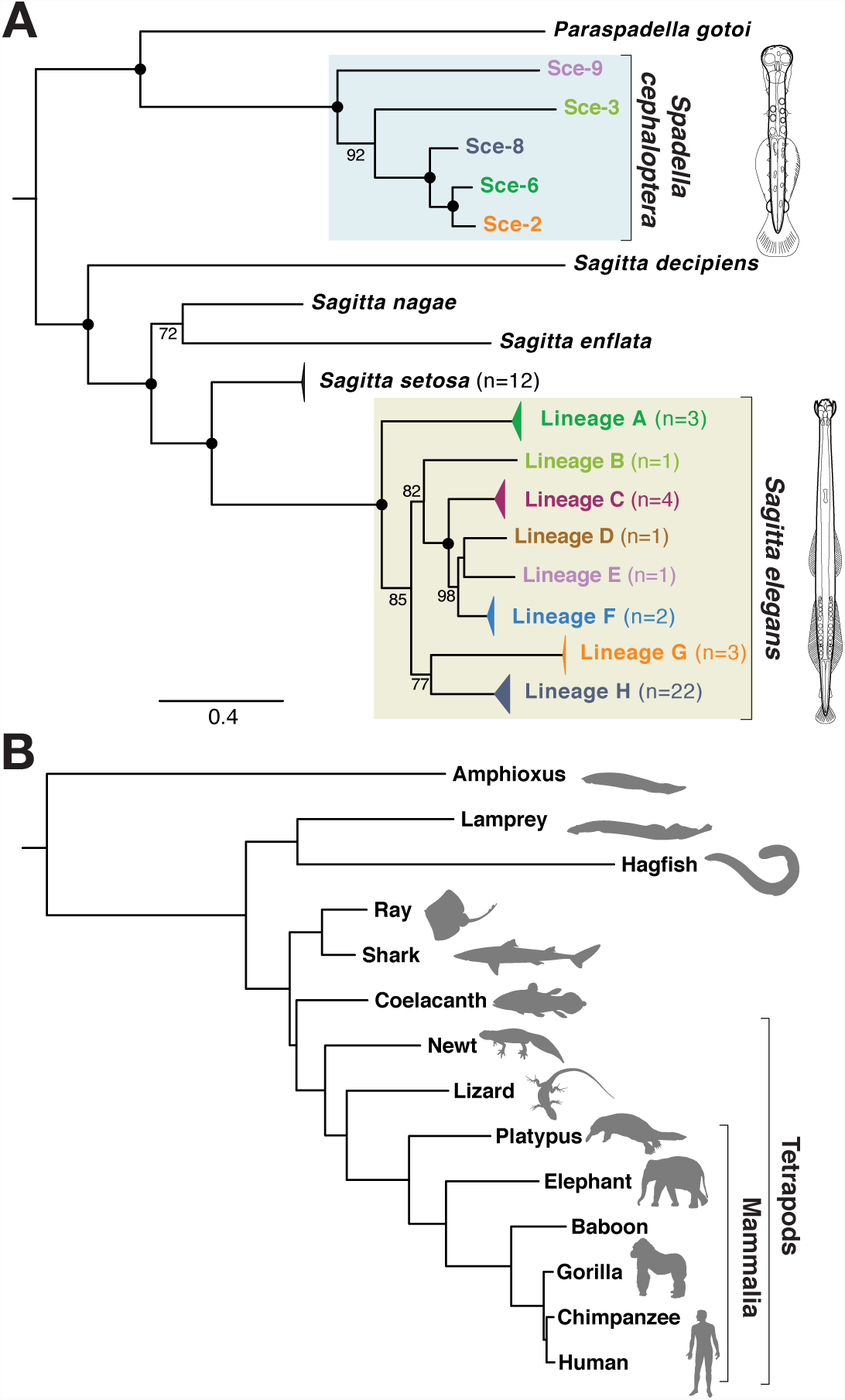
Mitochondrial divergence in chaetognaths and vertebrates. Phylogenetic trees based on the concatenation of all protein-coding mitochondrial genes in chaetognath lineages (A) and vertebrates (B) at the same scale (expected changes per site). Reconstructions were performed using Maximum Likelihood (MtZOA+r model). Maximal bootstrap support values are indicated by plain circles on nodes. In vertebrates, ML branch lengths were inferred from the alignment according to accepted topology. Sample sizes of *Sagitta elegans* lineages are indicated in brackets (detailed in S3 Figure).

### (d) Mitochondrial diversity values represent extremes across animals

We established that our nucleotide diversity estimates for two chaetognath species represent the highest values reported so far for any metazoan by surveying public databases for all available single-species population Cox1 datasets (Figure 3 and Table S2). *Sagitta elegans* shows the highest level of synonymous diversity for a single species (π_S_= 0.646) followed by *Spadella cephaloptera* (π_S_= 0.476). Conversely, *Spadella cephaloptera* is the more variable species at the coding level (π_N_= 0.023) followed by *Sagitta elegans* (π_N_= 0.018). We find that both chaetognaths harbor more intraspecific variation than several established cases of extreme mitochondrial diversity driven by (micro-) allopatric divergence, such as the crustacean *Tigriopus californicus* (π_S_= 0.404) and the gastropod *Cepaea nemoralis* (π_S_= 0.418) [30,31].

**Figure 3.**
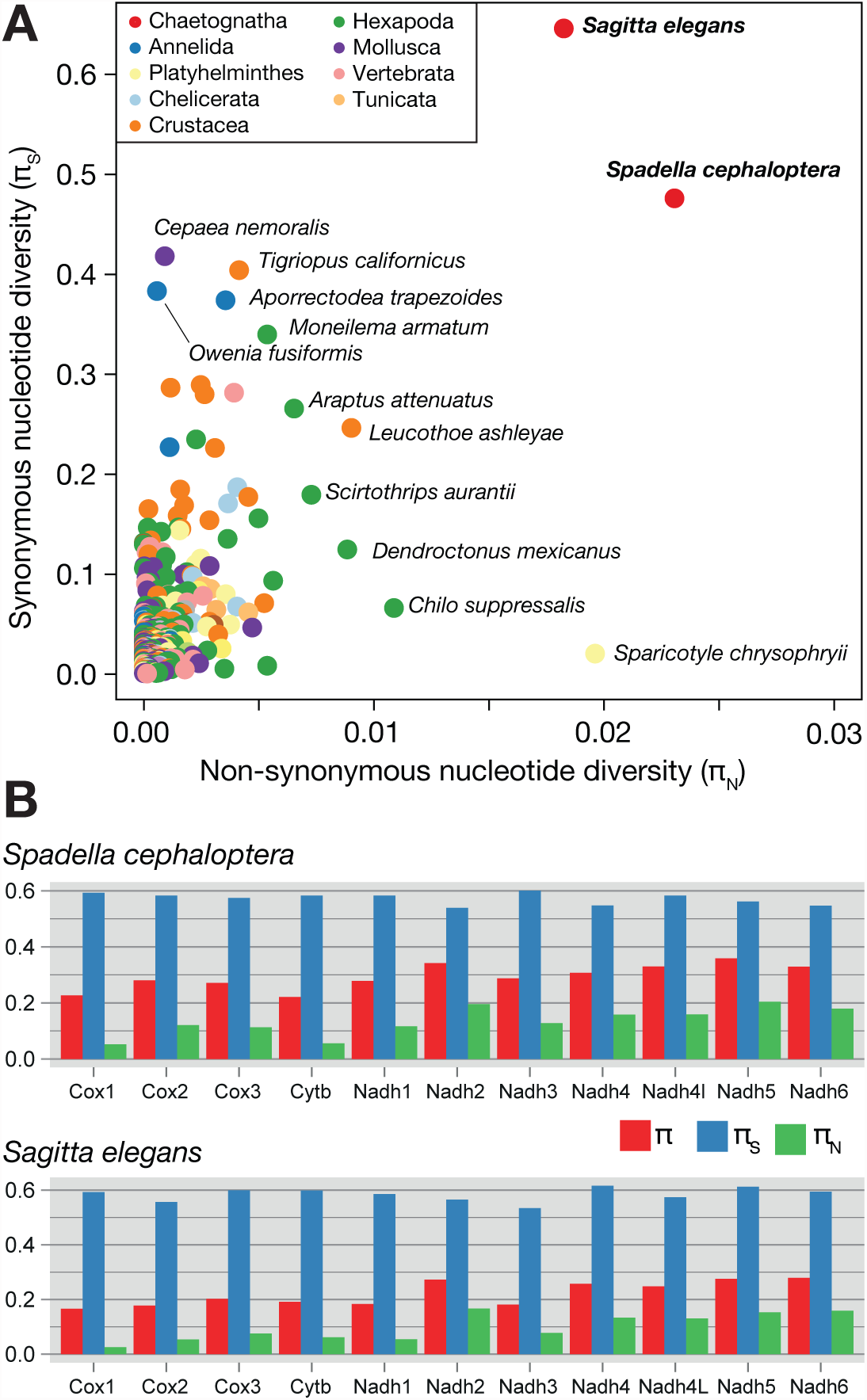
Mitochondrial diversity values in chaetognaths are highest among animals and homogenous across mitochondrial genes. (A) Nucleotide diversity computed for the 329 Cox1 datasets extracted from the NCBI popsets database that are associated with a referenced publication and do not constitute established cases of cryptic speciation. (B) Nucleotide diversities for all (π), non-coding (π_S_) and coding (π_N_) sites in 11 coding mitochondrial genes of *Spadella cephaloptera* (N =5) and *Sagitta elegans* (N=37).

### (e) Within-species mitochondrial rearrangements

*Sagitta elegans* mitochondrial lineages not only exhibit extreme molecular divergence, but also contain striking structural rearrangements (Figure 4). Changes in gene order are commonly observed between species [26,28,32], but to our knowledge such rearrangements have never been described as part of the allelic diversity of a single natural population before. In *Sagitta elegans*, at least two lineages exhibit rearrangements compared to the standard gene order involving the genes Nadh1, Nadh2 and Cox3 in one case, and Nadh4L in the other (Figure 4*a* and *4b*). Structural changes in mitogenomes also affect the size and proportion of intergenic regions, which are highly variable between lineages, ranging from 8.3% in lineage H to 25.4% in lineage G. Conversely, the fraction of non-coding DNA is remarkably stable within lineages with at most 3% variation (Figure 4*a*). These intergenic regions do not contain palindromic motifs which would be indicative of a repetitive element origin [33]. We also identified partial Cox1 extra-copies inserted in intergenic regions of *Spadella cephaloptera* mitogenomes Sce-2 and Sce-6, which had accumulated coding substitutions. These duplicates support the model that considers gene duplications as intermediate steps in structural rearrangements of the mitochondrion [34]. Such dynamic rearrangements could be responsible for the extreme reduction in size and gene number of chaetognath mitogenomes compared to other bilaterian animals, with the loss of ATP6, ATP8 and tRNA genes [27,28].

**Figure 4.**
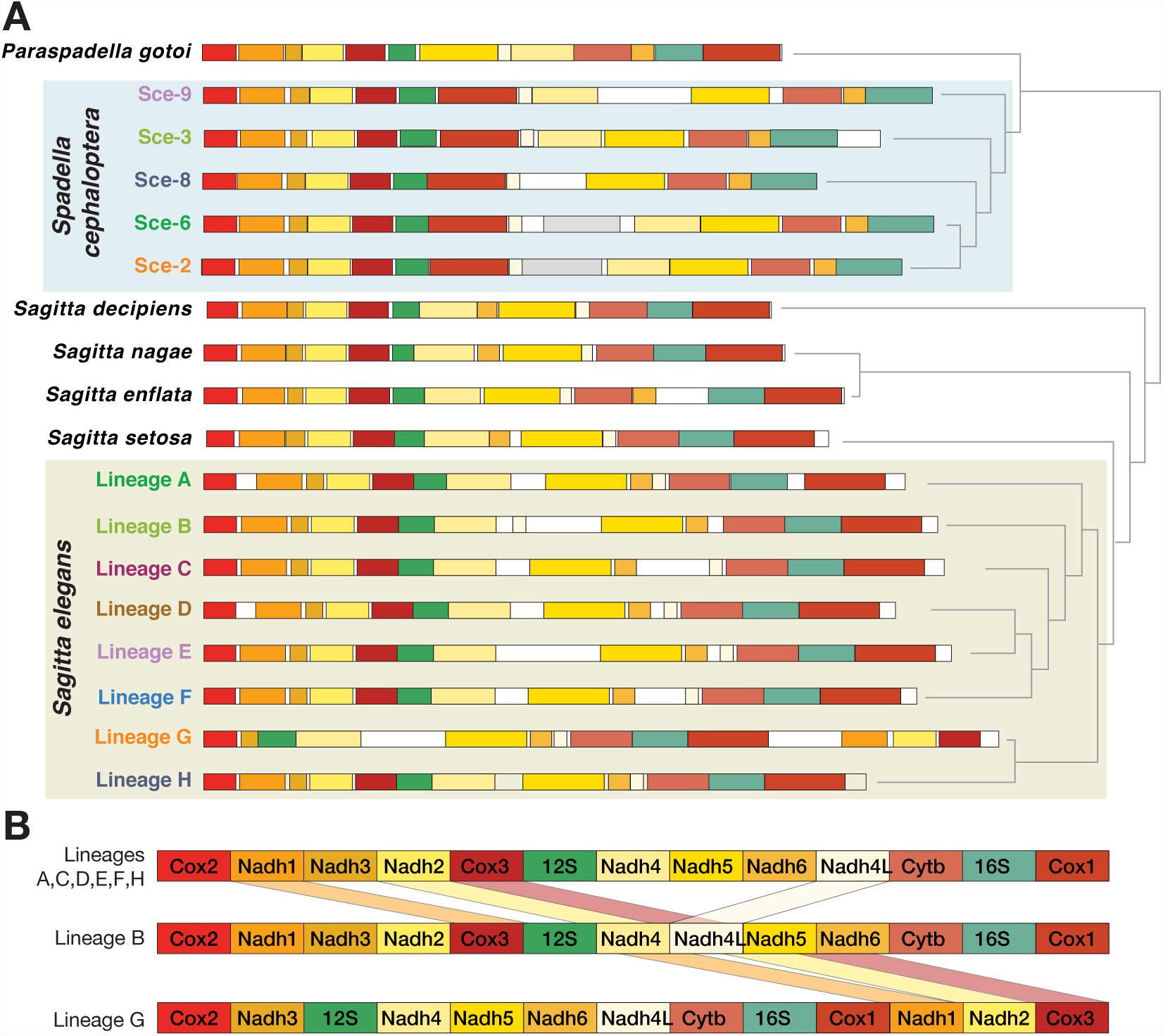
Structural variation between individual mitochondrial genomes of arrow worms. (A) Annotated mitochondrial genomes with gene positions drawn to scale. The proportion of intergenic regions (white) varies from 8% to 26.4% in *Sagitta elegans* lineages and from 3.7 to 18.4% in *Spadella cephaloptera* lineages. Cox1 duplicates in *S. cephaloptera* are in grey. (B) Schematic representation of putative mitochondrial gene order rearrangements in two *Sagitta elegans* mitochondrial lineages (B and G) compared to the most frequent gene order (lineages A,C,D,E,F,H).

### (f) Patterns of selection within mitogenomes

High levels of coding nucleotide diversity could be the result of positive selection [35]. Annotation of individual mitogenomes confirmed that the levels of nucleotide diversity are consistently high for all mitochondrial genes in *Spadella cephaloptera* and *Sagitta elegans* linages (Figure 3*b* and Table S3). We determined that remarkably high levels of synonymous diversity resulted in low or moderate π_N_/π_S_ ratios. Similar to what has been reported in mammals [36], subunits of the NADH complex exhibit higher non-synonymous variation than the subunits of the Cytochrome Oxidase complex suggesting shared evolutionary constraints across animals. To cope with mutational saturation in our data (Figure S5), we conducted maximum-likelihood estimates of d_N_/d_S_ ratios in *Sagitta elegans.* As we found low intraspecific pairwise d_N_/d_S_ ratio (0.056), we investigated whether individual lineages contribute unequally to the global d_N_/d_S_ estimates. We estimated independent d_N_/d_S_ ratios in each gene and lineage. This analysis does not support an increased d_N_/d_S_ in any particular lineage or gene, though some genes seem prone to higher coding variation, such as Nadh6 in Lineage G or Nadh3 in Lineage C (Figure S6). Similarly, likelihood-ratio tests did not provide significant support for positive selection affecting specific sites (Table S4). These analyses consistently indicate that positive selection is not responsible for the divergence of mitochondrial lineages in *Spadella cephaloptera* and *Sagitta elegans*, instead, the low d_N_/d_S_ values suggest that they have remained exposed to the influence of purifying selection during their evolution.

## 3. Discussion

We uncovered extreme mitochondrial diversity in single populations of the chaetognaths *Spadella cephaloptera* and *Sagitta elegans* and demonstrated that corresponding lineages are split by class-level molecular divergence and structural rearrangements. While cryptic speciation is a common explanation for such extreme mitochondrial divergence, we did not observe a pattern of divergence at any of the nuclear loci consistent with reproductive isolation. Moreover, the high number of observed mitochondrial lineages in both species (>8) would require a highly unlikely scenario of reproductive isolation followed by subsequent hybridization events. Moreover, this unlikely scenario would have to be independently repeated in both *S. cephaloptera* and *S. elegans* considered.

The mechanism by which highly divergent mitochondrial lineages have appeared and are maintained in these populations of arrow worms remains to be fully understood. By considering synonymous and non-synonymous rates of substitutions, we first ruled out that these lineages emerged through positive selection. The low or moderate d_N_/d_S_ ratios recovered instead are indicative of a large population size (Table S4) [8,13]. Since *Spadella cephaloptera* and *Sagitta elegans* are distantly related in the chaetognath tree (belonging to the orders Phragmophora and Aphragmorphora, respectively) and have very distinct ecologies (benthic and planktonic, respectively), this extreme mitochondrial diversity does not appear to be related to the population genetics of one particular species, and could represent a common condition in the phylum. Previous reports of high mitochondrial variation in other chaetognath species are consistent with this claim, even though these have generally been interpreted as evidence of cryptic speciation [37,38]. We did not however recover such highly divergent mitochondrial lineages in the species *Sagitta setosa*, which indicates that the presence of such lineages is not an obligatory condition in the phylum. The reduced variation compared to other chaetognaths has previously been related to the fragmented, coastal populations of *S. setosa*, which probably suffered severe population bottlenecks during recent glacial periods [39].

The presence of highly divergent mitochondrial lineages therefore fits theoretical expectations predicting that a large population size allows for the maintenance of ancestral polymorphisms [6,7]. Hence, chaetognath mitochondrial diversity may be an exception to the ‘paradox of variation’ as our results challenge previous reports suggesting that the extent of mitochondrial variation is limited [8]. As pointed out by our molecular dating analyses, an ancient origin combined with fast mutation rates probably contributed to the evolution of divergent mitochondrial lineages in chaetognaths (Figure S4). Such accelerated mutation rates could be related to the extreme reduction of chaetognath mitogenomes and their propensity to structural rearrangements [26]. Such a combination of extreme size reduction and accelerated mutation has also been observed in ctenophores [40]. The size reduction and peculiar architecture of animal mitochondrial genomes compared to other eukaryotes, such as green plants, has previously been attributed to the combined effect of genetic drift and accelerated mutation rates [41]. The processes that gave rise to the extreme mitochondrial diversity in chaetognaths could therefore be similar to those governing mitochondrial evolution in other metazoans, but only differ by the extent of mutation rate acceleration and population size.

Extreme levels of intraspecific mitochondrial divergence also imply that cyto-nuclear interactions at the respiratory complexes are much less constrained than originally thought [11]. Indeed, the interactions between mitochondrial and nuclear subunits should be robust enough to cope with multiple divergent mitochondrial lineages present in interbreeding populations of chaetognaths. Mitochondrial transplantation experiments originally demonstrated that cytonuclear interactions could be maintained between species, but were dramatically altered when divergence increased (*e.g.* to human and orang-utan), resulting in a decreased efficiency of respiratory processes. Similar effects were reported in hybrids between divergent *Tigriopus californicus* populations, which showed reduced respiratory fitness and altered gene expression [12,42]. We hypothesize that chaetognaths cope with their high levels of mitochondrial divergence through an increased robustness in the interaction with nuclear subunits of oxidative phosphorylation complexes. Such a higher robustness could be achieved by adaptive amino acid changes in nuclear subunits or could derive from modifications in the respiratory subunit gene repertoire, for instance through gene duplication, a frequent process in chaetognaths [16].

An important problem remains to determine whether chaetognaths constitute a unique case or whether extreme mitochondrial diversity is more common in animals. Many instances of high mitochondrial diversity have directly or indirectly been interpreted as evidence of cryptic speciation (for instance, see [43-46]). Some of these cases may be subject to reevaluation when investigated using nuclear loci. Alternatively, other reasons why similar cases of high mitochondrial diversity could have been overlooked may be that sequences from highly divergent individuals are simply not amplified through PCR-based approaches because of mismatches in the primer regions, or that strongly divergent sequences are discarded as contaminations or misidentifications. Our findings therefore prompt a reassessment of the classical DNA barcoding workflow, suggesting in particular that mitochondrial markers should not be considered without nuclear counterparts, and that extra attention should be paid to outliers and unsuccessful PCR amplifications.

Our results challenge established views of the amount of mitochondrial diversity that can be harboured within single species. An assessment of intraspecific mitochondrial diversity in other animal groups, particularly those with dynamic mitochondrial genomes, such as ctenophores, urochordates or brachiopods, may uncover other exceptions [6]. Until more of these hyper-diverse taxa are identified, chaetognaths represent a new and pivotal model to understand the molecular evolution of animal mitochondrial genomes.

## 4. Materials and methods

### (a) Animal sampling and genotyping

Individuals of the species *Spadella cephaloptera*, were collected from Calanque de Sormiou near Marseille (France) at shallow depth on *Posidonia* seagrass using a plankton net (Figure S1). Individuals of the species *Sagitta elegans* and *Sagitta setosa* were collected using a multinet on a single daytrip in the Gullmar fjord near Kristineberg (Lysekil, Sweden). All chaetognaths were identified under a microscope while still alive. Only individuals without food in their guts and without visible parasites were preserved and used for genetic analysis.

Genomic DNA was extracted using Qiamp Micro Kit and DNeasy Blood & Tissue Kit (Qiagen) from 30 *Spadella cephaloptera*, 109 *Sagitta elegans* and 57 *Sagitta setosa* individuals and used as a template for PCR amplification using GoTaq (Promega) or Phire Hot Start II DNA Polymerase (Finnzymes). For *Spadella cephaloptera*, fragments of the Cox1 and 16S mitochondrial genes were amplified for 25 and 30 individuals, respectively (Table S1). Fragments of Cox1 and Cox2 genes were recovered for 107 and 108 individuals of *Sagitta elegans* and 54 individuals in *setosa* (Table S1).

A fragment spanning the nuclear intron of the ribosomal protein L36a gene was amplified in 30 *Spadella cephaloptera* and the 37 *Sagitta elegans* individuals including all the individuals for which the entire mitogenome was sequenced. The primers employed were designed to specifically amplify L36a, one of the two ancient chaetognath specific duplicates of the L36 genes detected in *S. cephaloptera* [47]. Furthemore, for *S. elegans* and *S. setosa* fragments of 18S and 28S ribosomal RNA, and for *Spadella cephaloptera* of 18S ribosomal RNA were amplified using primers and protocols described in ref. [23]. All amplified fragments were sequenced using Sanger technology in both directions, and for the L36a intron of both species, and 18S and mitochondrial genes of *Spadella cephaloptera*, the fragments were cloned in pGEM-T easy (Promega) prior to sequencing. All primers employed are specified in Table S1. All sequences were deposited in Genbank (see Table S1 for accessions).

### (b) Sequencing of individual mitochondrial genomes

Long-range PCR amplifications of whole mitochondrial genomes were carried out for all three chaetognaths *Spadella cephaloptera*, *Sagitta elegans* and *S. setosa.*

In *Spadella cephaloptera*, two half-genome fragments (6–8kb) were amplified using specific primers located in 16S and Cox1 genes using the Accuprime Taq (Invitrogen) for five selected individuals. These fragments were purified using S.N.A.P. gel purification kit (Invitrogen) and subsequently cloned with Topo XL cloning Kit (Invitrogen). Plasmid DNA was prepared with Plasmid Midi Kit (Qiagen). These templates were sequenced using a primer walking strategy: 6 to 10 Sanger reads were iteratively carried out using custom designed primers from both extremities of the insert using a primer walking strategy using 6 to 10 Sanger reads in total.

In *Sagitta elegans* and *Sagitta setosa*, fragments spanning the entire mitochondrial genome were obtained for 37 and 12 individuals, respectively. The *S. elegans* individuals are representative of eight deep mitochondral lineages recovered in that species. Outward pointing primers were designed within the Cox1 or the Cox2 genes and used to amplify the mitogenome as a single amplicon using Phusion DNA polymerase (Thermo scientific) with cycling parameters adjusted for long products. PCR products were purified using QIAquick purification kit (Qiagen). These products were processed for shotgun sequencing using a Next Generation Sequencing approach at the Wellcome Trust Center for Human Genetics (Oxford). A unique library was built for each individual mitogenome using Nextera Kit (Illumina) from 30-70 ng of PCR product. Multiplexed libraries were then sequenced on a single Illumina MiSeq run. An average of 71,000 paired-end reads (150bp) was obtained for each fragment library allowing for a median 800× coverage per genome. After de-multiplexing, reads from each library were trimmed for low quality stretches using sickle (available at github.com/najoshi/sickle), then error-corrected and assembled with SPAdes (v2.5.0) using BayesHammer error correction and multiple k-mers (21,33,55,77) [48]. To check for potential mis-assemblies, reads were mapped back to each mitochondrial genome using Bowtie2 [49] and corresponding alignments were inspected by eye in IGV viewer [50] to check even coverage and concordance of pair-end information with the assembled sequence. The scaffold with highest coverage was selected and compared with Cox1 and Cox2 fragments previously obtained. This procedure ascertained that no sample mixing happened during the sequencing process and confirmed the identity of each individual mitochondrial genome.

### (c) Annotation of mitochondrial genomes

For all mitogenomes, ORFs from all-three frames were searched for protein coding genes by tblastn using available chaetognath sequences as queries. Corresponding sequences were extracted, translated and aligned. Inspection of amino acid alignments and comparison with predictions available for other chaetognath species helped to define start codons and gene boundaries. In-frame nucleotide alignments were constructed from the validated amino acid analyses and subsequently used in further molecular evolution analyses. Similarly, non-coding genes (12S and 16S rRNA) were extracted based on similarity and gene borders defined using multiple alignments. Refined amino acid and nucleotide sequences were aligned back to each genome for annotation and determination of gene order (Figure 4).

### (d) Molecular evolution and population genetic analysis

Standard population genetic parameters were calculated from nucleotide datasets using the EggLib python library [51]. To show evolutionary relationships of mitochondrial haplotypes, we used Neighbor-Net Networks from Splitstree4 with K2P distances [52] (Figure 1 and Figure S2). We also performed Maximum Likelihood (ML) phylogenetic inferences using RAxML assuming the GTR+Г model for nucleotide datasets and the MtZoa+ Г model for amino acid datasets [53,54] (Figure 2*a*). Saturation levels in each marker were evaluated for both amino acid and nucleotide data by comparison of pairwise distances under a simplistic model of evolution (Kimura protein, K2P nucleotide) and the patristic ML distance (GTR+Г or MtZOA+Г).

Non-synonymous and synonymous substitution rates were estimated following distinct approaches using the PAML package [55]: (i) pairwise comparisons were conducted on each gene using the yn00 program [56], (ii) to account for potential lineage specific effects, the ‘branch-model’ was employed in codeml assuming a different d_N_/d_S_ ratio in each lineage for *S. elegans* (Figure S6), and (iii) the ‘site-model’ in codeml was applied to evaluate selective effects at specific codons (Table S4).

### (e) Survey of mitochondrial genetic diversity in metazoans

To explore the extent of mitochondrial genetic variation in metazoans, we searched the NCBI popset database for datasets corresponding to Cox1 gene fragments (keyword: Cox1, COI, CO1), including a single species and at least 10 sequences. We obtained 1581 records corresponding to 1079 species. We focused on the 437 datasets associated with a referenced publication (pmid) and we excluded datasets corresponding to cryptic species based on keyword search in abstract of papers (keywords: ‘cryptic’ and ‘species complex’), and manual curation, yielding 329 datasets. We retrieved sequences from the corresponding datasets, and built a codon alignment based on the alignment of amino acid translation using Muscle. Distinct fragments of Cox1 were sometimes pooled in the same popset causing misalignment, in which cases we discarded the shorter fragments. We also discarded sequences including degenerate bases (e.g. ‘N’, ‘Y’ or ‘R’). We computed nucleotide diversity statistics using the egg-lib library [51] and we selected the dataset showing the highest diversity in each species (Table S2). We then verified the alignment and corresponding papers in the 50 datasets with highest levels of genetic diversity.

### (f) Molecular dating analysis

To approximate divergence times and evolutionary rates, we conducted a molecular dating analysis using a concatenated amino acid alignment of all mitochondrial genes for both chaetognaths and several representative vertebrate species (Figure S4). We set calibration points for vertebrates according to [57]. For chaetognaths, we specified the divergence of Aphragmophora and Phragmophora to range between 530 Myr and 250 Myr. 530 Myr corresponds to the age of the first unquestionable chaetognath fossil [58], and 250 Myr represents the onset of Tethys ocean opening [59]. Given that this geological event matches the geographic distribution of several chaetognath species belonging to either Aphragmophora or Phragmophora, including *Spadella cephaloptera,* the divergence between the two classes most likely predates this event. Similarly, we considered that the multiple lineages in *S. cephaloptera* must have diverged after 250 Myr because of the tethysian distribution of this species. Phylobayes was employed for molecular dating assuming a log-normal autocorrelated relaxed clock and a prior of 700 Myr with 200 Myr deviation on the age of the root [60]. Chains were run for at least 20,000 cycles and 5,000 were discarded as burn-in.

## 6. Acknowledgements

This research was supported by NWO-VENI grant 863.08.024 to K.P. and by ERC (FP7/2007– 2013)/ERC grant [268513]11 funding F.M. The Wellcome Trust Centre for Human Genetics (WTCHG) is funded by Wellcome Trust grant 090532/Z/09/Z and MRC Hub grant G0900747 91070.

We thank L. Friis Møller and C. Hörnlein for help with sampling in Gulmar fjord in Sweden; E. Goetze, M. Taylor, L.E. Becking, I. Maeso, J. Paps, W. Renema, M. Schilthuizen and P. W. H. Holland for commenting on the manuscript; E.J. Bosch for help with illustrations; P. Kuperus and C. Ribout for technical assistance and the WTCHG for the generation of the sequencing data.

## Author contribution

F.M., Y.L.P. and K.P. conceived the project and carried out the animal sampling. F.M. and S.L. performed the experiments. F.M., S.L. and Y.L.P. analysed the sequencing data. F.M. and K.P. wrote the paper.

## Data accessibility

- Assembled mitochondrial genomes have been deposited in Genbank under the accessions KP899748-KP899801 (details in Table S1)
- Sequences of genotyped markers are available under the accessions: KP843748-KP843841 and KP857119-KP857568 (details in Table S1).
- Genotyped markers, alignments, and reconstructed trees are also available as Dataset S1 and S2 at http://dx.doi.org/10.5061/dryad.66hg0.

---

**Figure S1.**
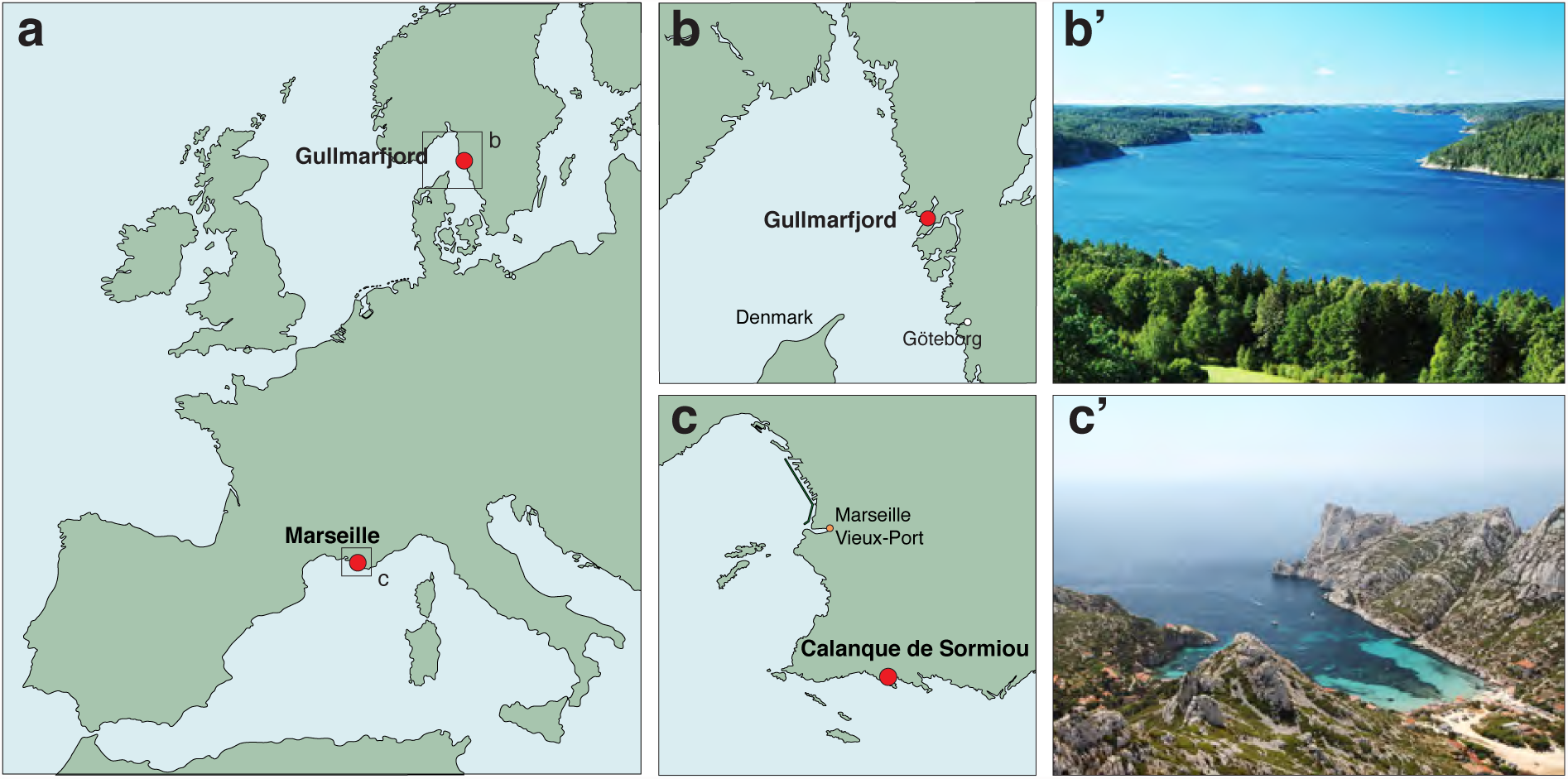
Sampling locations for *Spadella cephaloptera* (Calanque de Sormiou, France) and *Sagitta elegans* and *S. setosa* (Gullmar fjord, Sweden).

**Figure S2.**
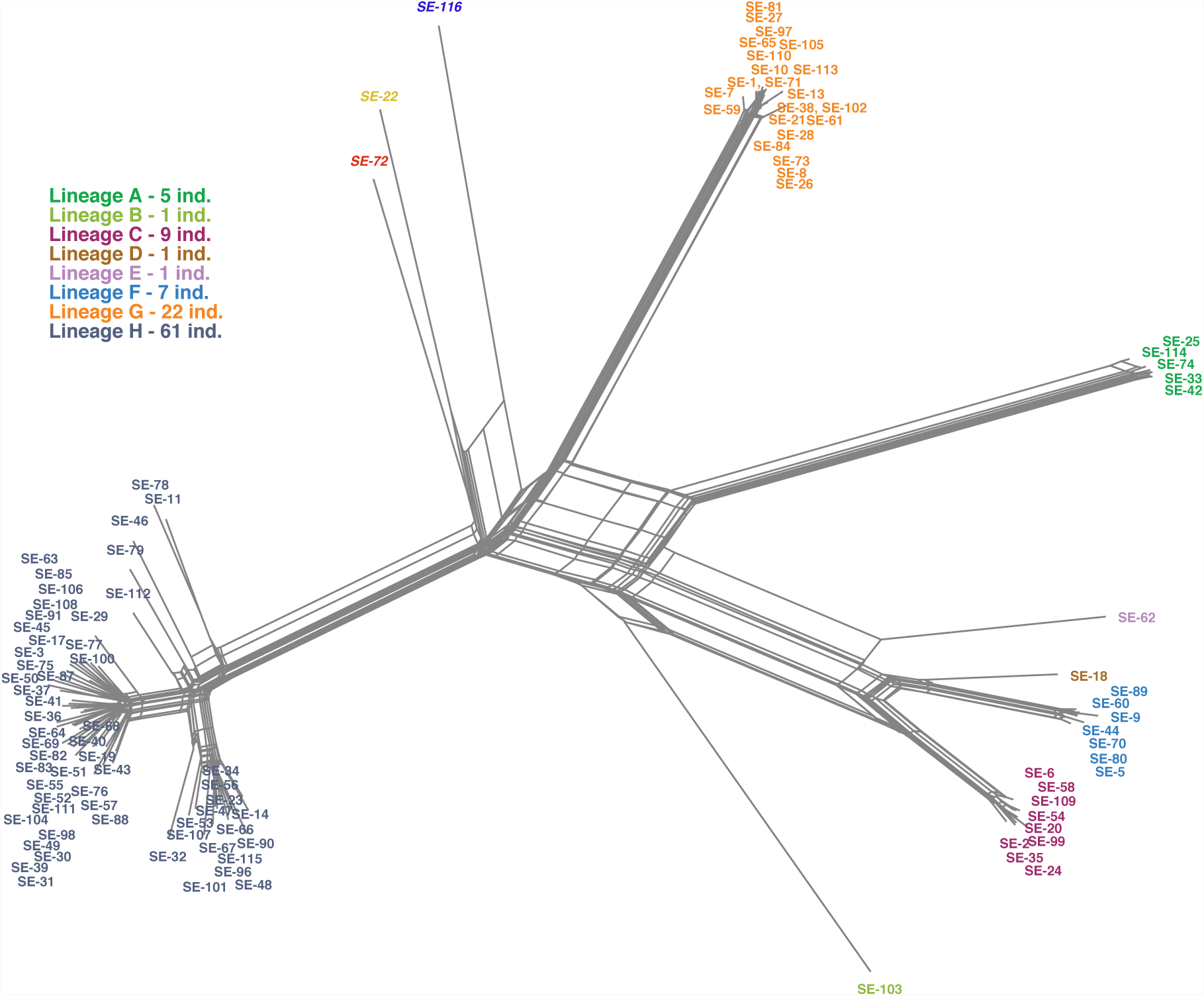
Neighbor-net phylogenetic network constructed using K2P distances between Cox1 sequences from 107 *Sagitta elegans* individuals randomly sampled from Gullmar fjord. Individual numbering is colored according to lineage assignment.

**Figure S3.**
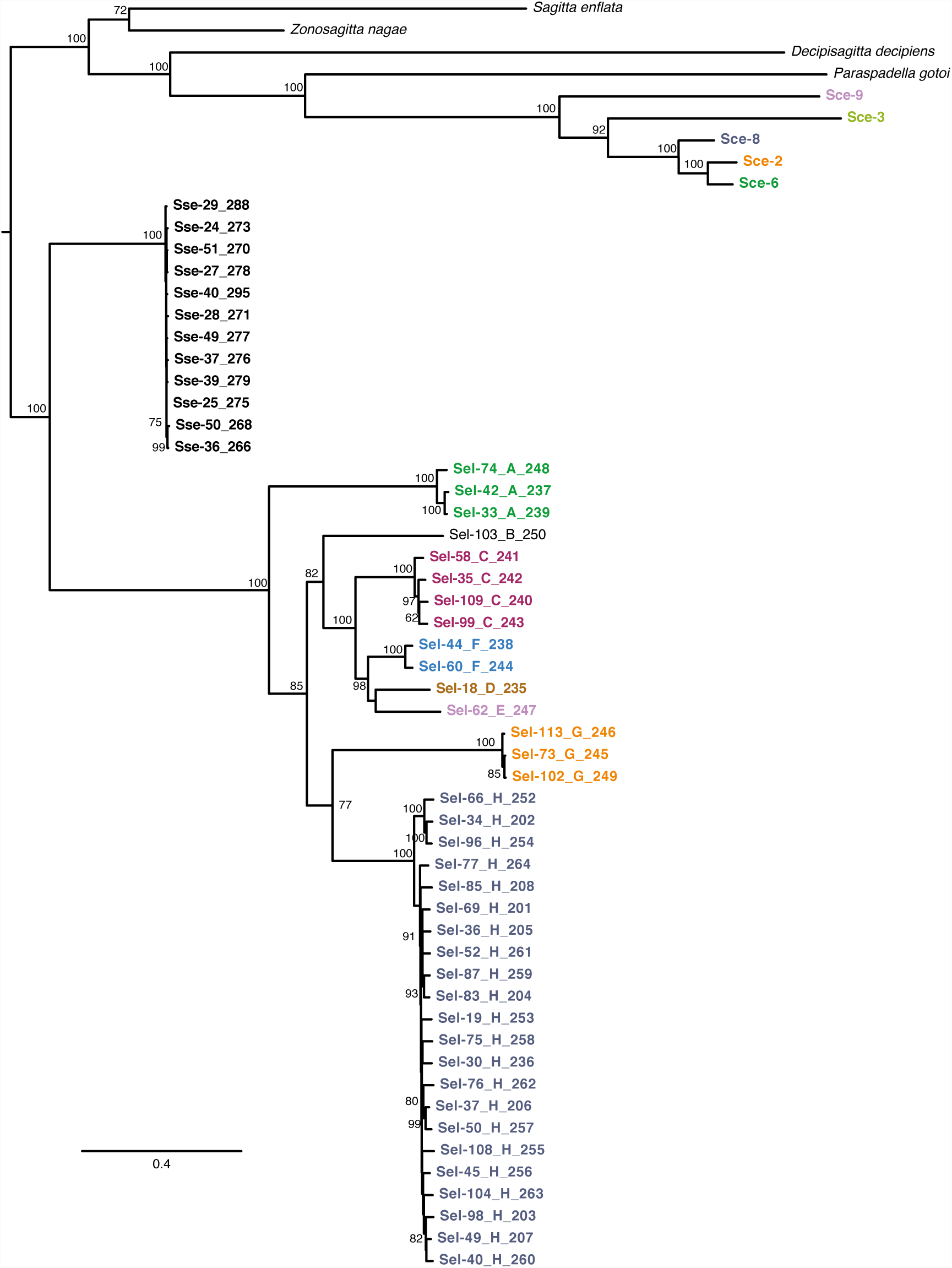
Maximum-likelihood tree of concatenated protein-coding mitochondrial genes in chaetognaths including all newly sequenced mitochondrial genomes for *Spadella cephaloptera* (Sce), *Sagitta elegans* (Sel), and *Sagitta setosa* (Sse). Branch support was estimated from 400 bootstrap replicates (determined by autoMRE criterion) and only included if above 50.

**Figure S4.**
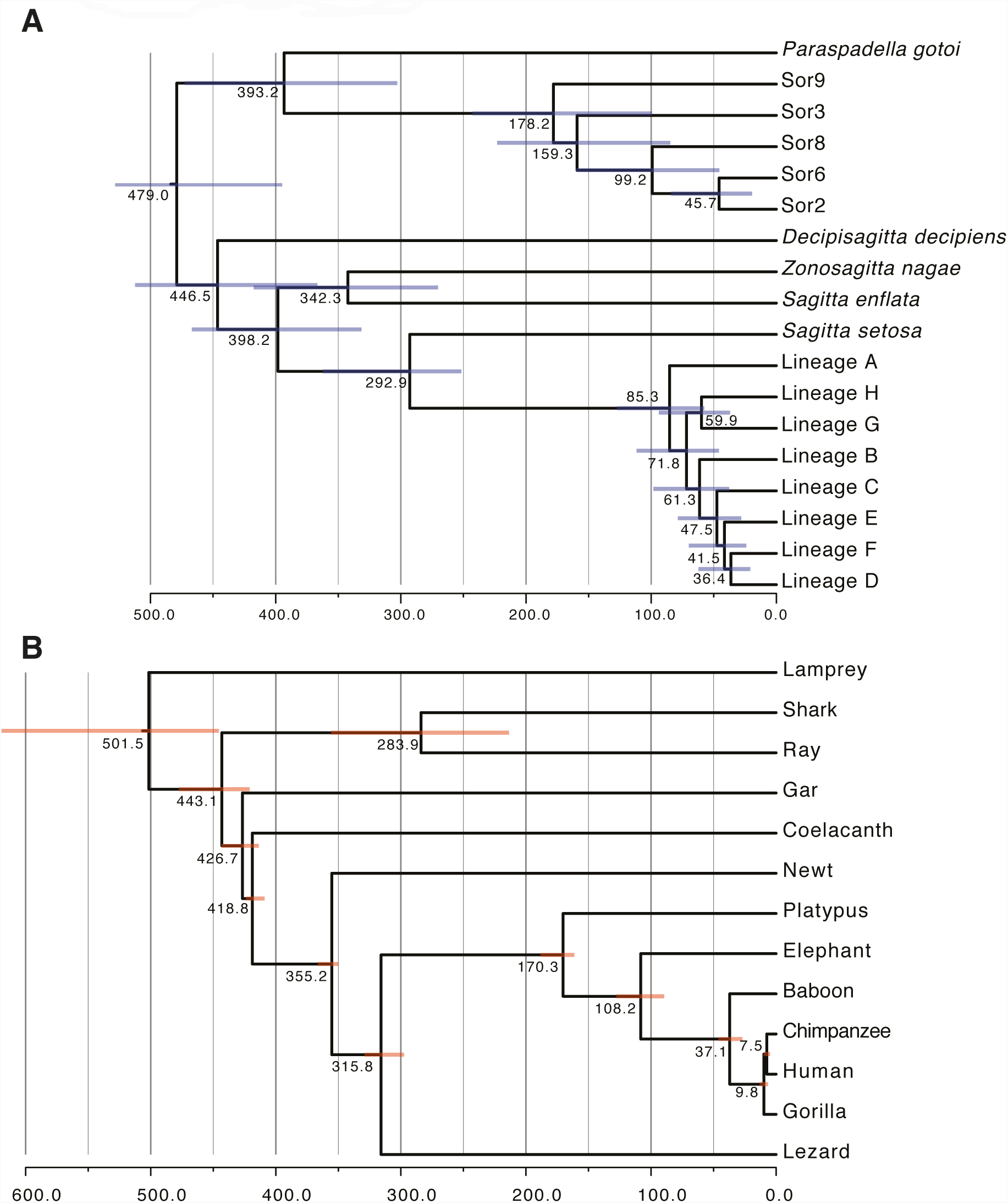
Molecular dating of chaetognath lineages in comparison with vertebrates. Chronograms derived from Phylobayes analysis are shown on the same scale, which represents million years divergence since present for (A) chaetognaths and (B) vertebrates. Error bars at nodes indicate the 95% Bayesian credibility interval estimated from each dataset.

**Figure S5.**
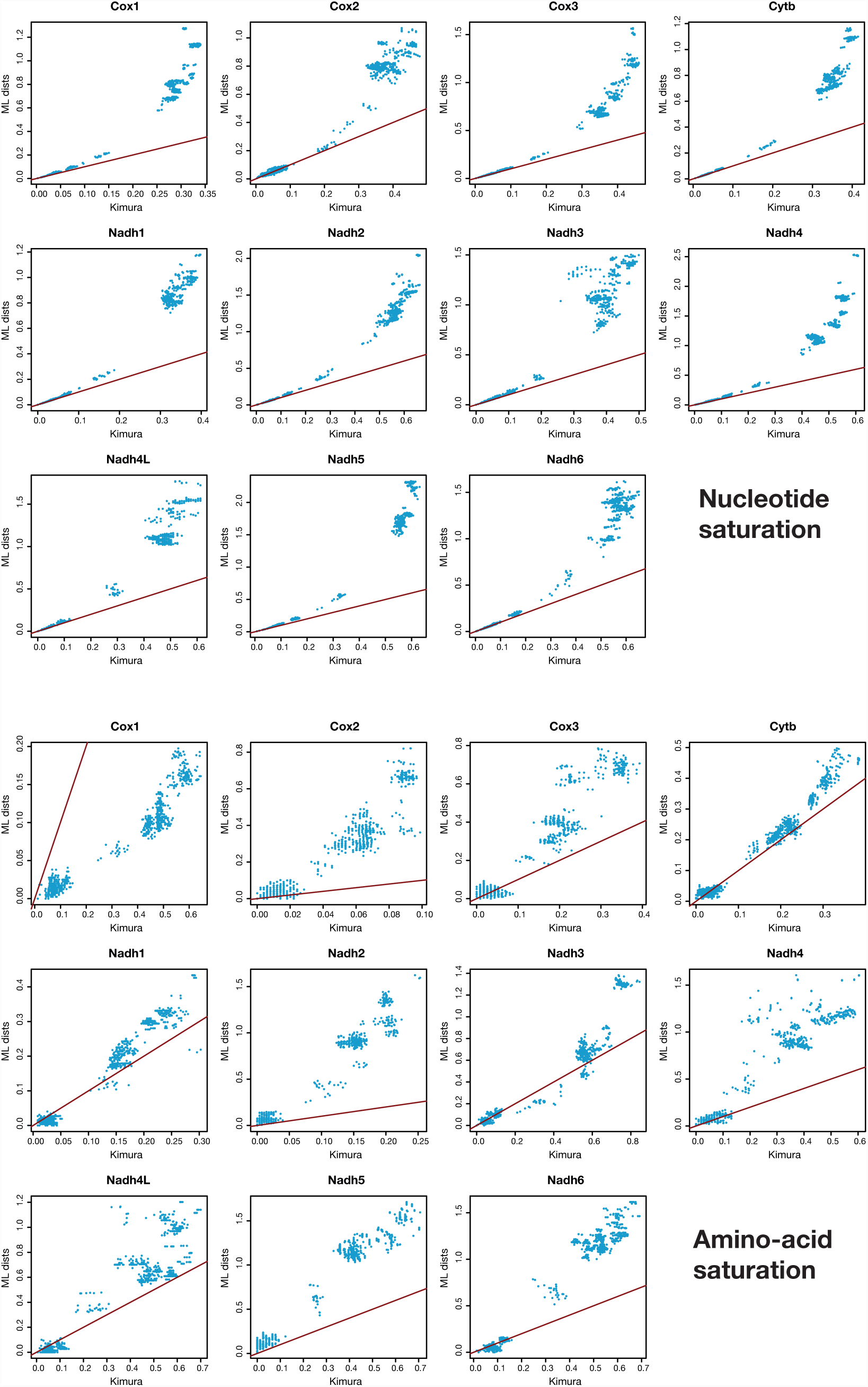
Saturation analysis in *Sagitta elegans* mitochondrial genes. Pairwise distances (K2P or Kimura protein distance) were compared with patristic ML distances for each marker. Deviation from the diagonal (dark red) is an indication of the degree of mutational saturation in a dataset. For instance, Cytb or Cox1 amino-acid datasets are less saturated than Nadh5 and Nadh6.

**Figure S6.**
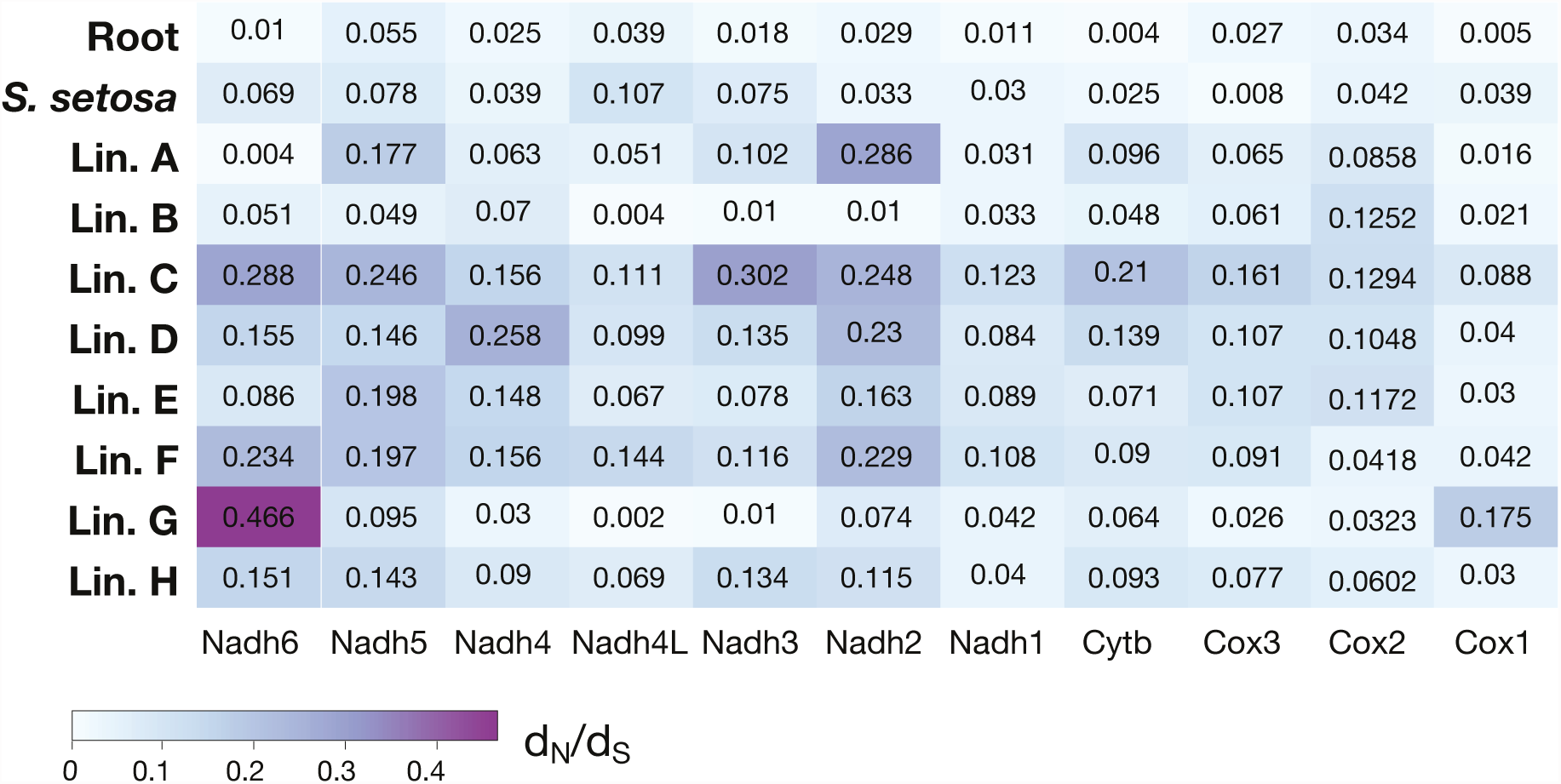
Heatmap representation of per-lineage d_N_/d_S_ estimates using PAML ‘branch-model’ in each mitochondrial gene of *Sagitta elegans*. Distinct d_N_/d_S_ ratios were assigned for each gene to the internal branches of the tree (‘base’) leading to *Sagitta setosa* and to the multiple *Sagitta elegans* deep lineages.

**Table S1.** Genotyped markers and corresponding summary statistics for single populations of *Spadella cephaloptera*, *Sagitta elegans*, and *Sagitta setosa*.

| Marker | nind | nseff | lseff | K | S | $\eta$ | $\theta_w$ | $\pi$ | $D_T$ | Accession |
| --- | --- | --- | --- | --- | --- | --- | --- | --- | --- | --- |
| *Spadella cephaloptera* |  |  |  |  |  |  |  |  |  |  |
| Cox1 | 25 | 25 | 639 | 25 | 295 | 439 | 0.1223 | 0.1385 | 0.5319 | KP843778-KP843802 |
| 16S | 30 | 30 | 549 | 29 | 250 | 359 | 0.1149 | 0.1181 | 0.1066 | KP843748-KP843777 |
| L36a intron | 30* | 39 | 652 | 36 | 122 | 132 | 0.0443 | 0.0411 | -0.2605 | KP843803-KP843841 |
| 18S | 8* | 8 | 1021 | 8 | 27 | 27 | 0.0102 | 0.0095 | -0.3362 | KP857143-KP857150 |
| mitogenomes | 5* |  |  |  |  |  |  |  |  | KP899748-KP899752 |
| *Sagitta elegans* |  |  |  |  |  |  |  |  |  |  |
| Cox1 | 107 | 107 | 578 | 96 | 259 | 494 | 0.0854 | 0.1775 | 3.6069 | KP857408-KP857514 |
| Cox2 | 108 | 108 | 376 | 101 | 220 | 432 | 0.1113 | 0.2043 | 2.7856 | KP857246-KP857353 |
| Cox1 full-length | 37* | 37 | 1521 | 37 | 646 | 1126 | 0.1017 | 0.1663 | 2.4035 | see mitogenomes |
| L36a intron | 37* | 71 | 345 | 70 | 98 | 109 | 0.0588 | 0.0276 | -1.8070 | KP857178-KP857245 |
| 18S | 24* | 24 | 1760 | 2 | 0 | 0 | 0.0000 | 0.0000 | NA | KP857119-KP857142 |
| 28S | 27* | 27 | 942 | 1 | 0 | 0 | 0.0000 | 0.0000 | NA | KP857151-KP857177 |
| mitogenomes | 37* |  |  |  |  |  |  |  |  | KP899765-KP899801 |
| *Sagitta setosa* |  |  |  |  |  |  |  |  |  |  |
| Cox1 | 54 | 54 | 655 | 49 | 64 | 69 | 0.0214 | 0.0088 | -2.0446 | KP857515-KP857568 |
| Cox2 | 54 | 54 | 504 | 53 | 70 | 76 | 0.0305 | 0.0138 | -1.9015 | KP857354-KP857407 |
| Cox1 full-length | 12* | 12 | 1521 | 10 | 46 | 50 | 0.0107 | 0.0079 | -1.2842 | see mitogenomes |
| mitogenomes | 12* |  |  |  |  |  |  |  |  | KP899753-KP899764 |

Cox1: Cytochrome Oxidase 1, Cox2: Cytochrome Oxidase 2, 16S : 16S mitochondrial ribosomal DNA, L36a: L36a nuclear intron, 18S: small subunit nuclear ribosomal DNA, 28S: large subunit nuclear ribosomal DNA. nind: number of individuals, nseff : number of sequences, lseff : number of analysed sites, K: number of haplotypes, S: number of polymorphic sites, $\eta$ : minimal number of mutations, $\theta_w$ : Watterson estimator, $\pi$ : nucleotide diversity, $D_T$ : Tajima's D; \* non-random sampling as individuals were selected to represent different mitochondrial lineages.

**Table S2.** Population genetic statistics calculated for each single-species NCBI ‘popset’ and used for Fig. 2. (see excel file).

**S3 Table.**
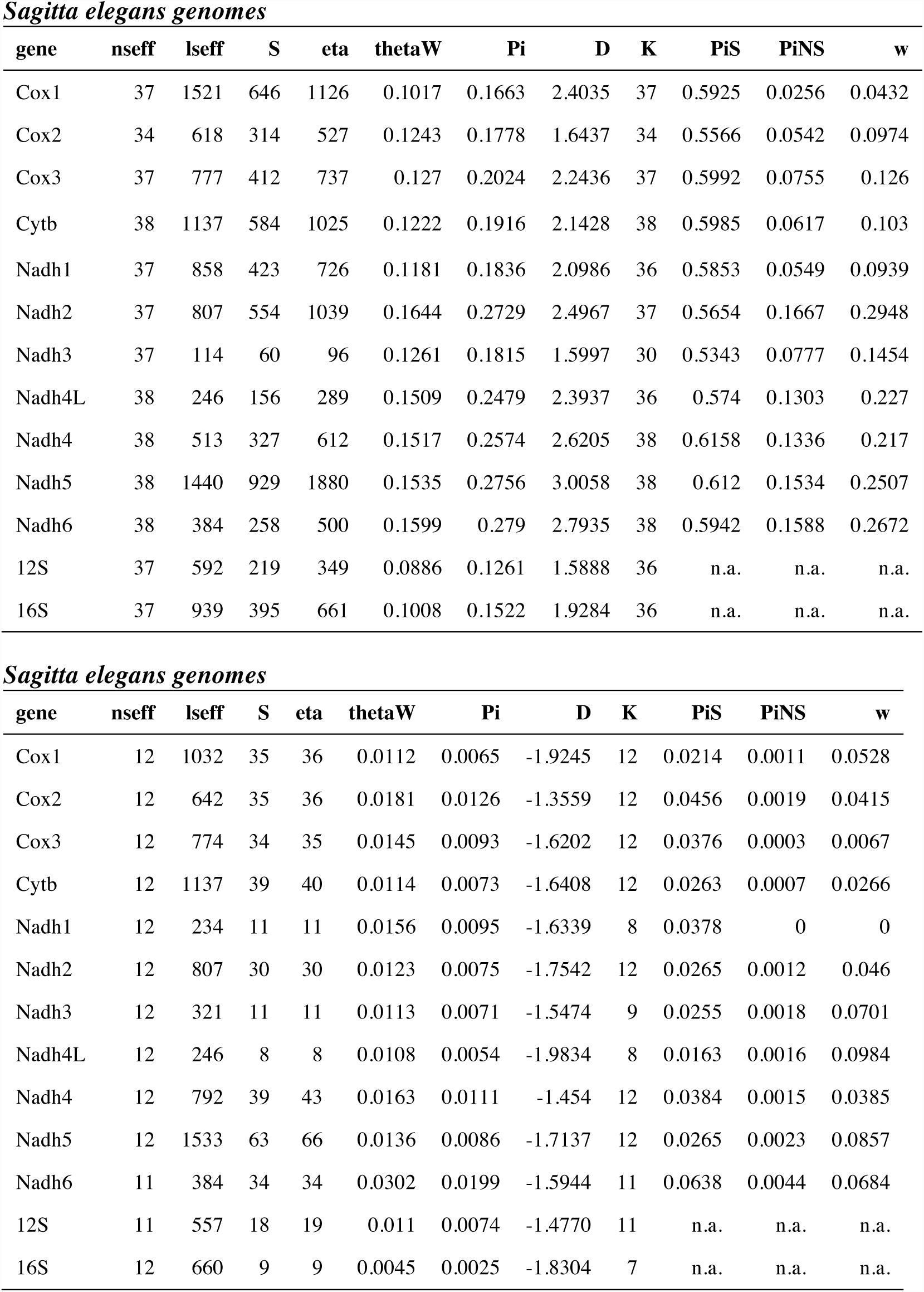
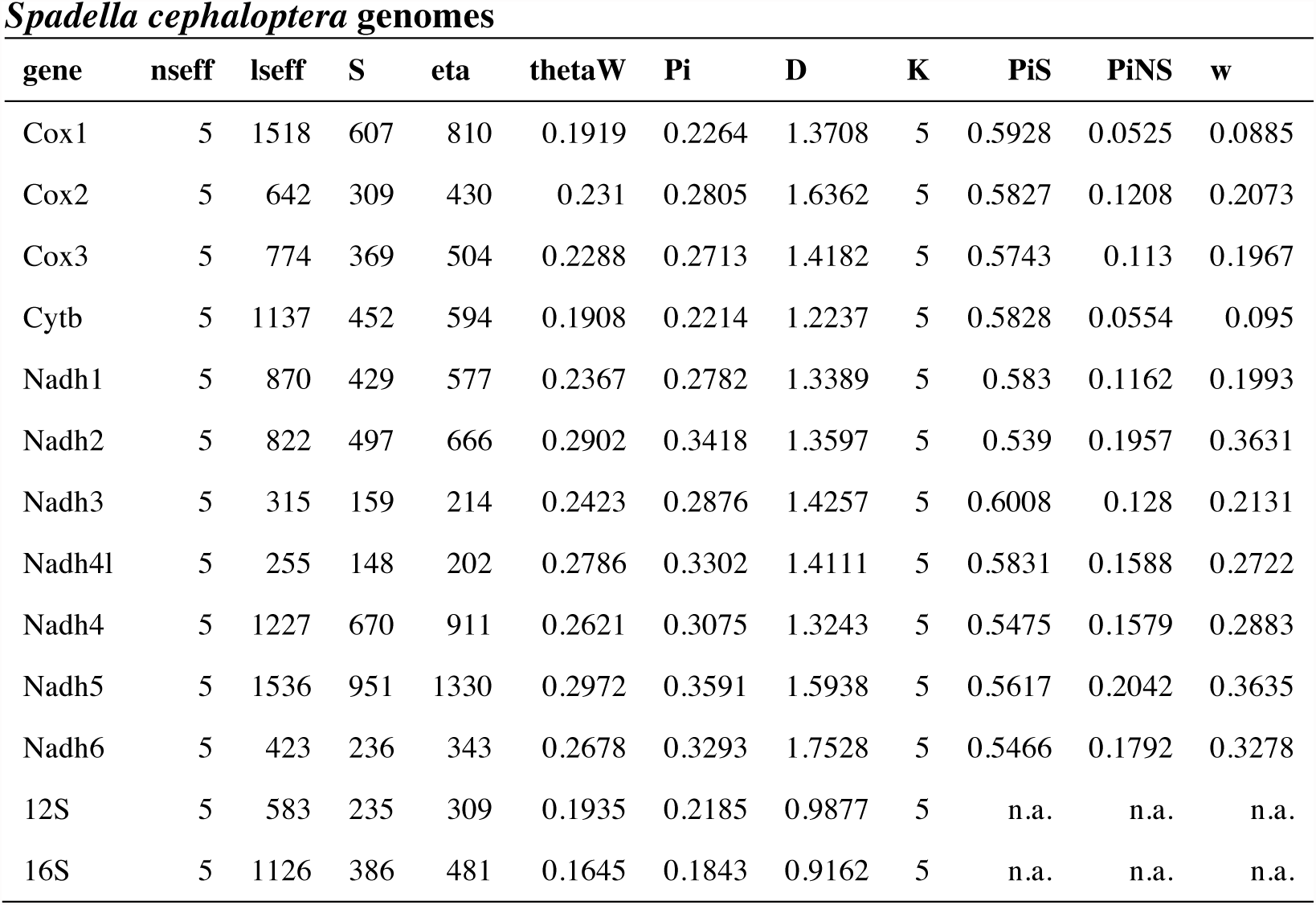
Population genetic statistics calculated for each mitochondrial gene in sequenced mitogenomes of *Sagitta elegans*, *S. setosa* and *Spadella cephaloptera*.

| gene | nseff | lseff | S | eta | thetaW | Pi | D | K | PiS | PiNS | w |
| --- | --- | --- | --- | --- | --- | --- | --- | --- | --- | --- | --- |
| Cox1 | 37 | 1521 | 646 | 1126 | 0.1017 | 0.1663 | 2.4035 | 37 | 0.5925 | 0.0256 | 0.0432 |
| Cox2 | 34 | 618 | 314 | 527 | 0.1243 | 0.1778 | 1.6437 | 34 | 0.5566 | 0.0542 | 0.0974 |
| Cox3 | 37 | 777 | 412 | 737 | 0.127 | 0.2024 | 2.2436 | 37 | 0.5992 | 0.0755 | 0.126 |
| Cytb | 38 | 1137 | 584 | 1025 | 0.1222 | 0.1916 | 2.1428 | 38 | 0.5985 | 0.0617 | 0.103 |
| Nadh1 | 37 | 858 | 423 | 726 | 0.1181 | 0.1836 | 2.0986 | 36 | 0.5853 | 0.0549 | 0.0939 |
| Nadh2 | 37 | 807 | 554 | 1039 | 0.1644 | 0.2729 | 2.4967 | 37 | 0.5654 | 0.1667 | 0.2948 |
| Nadh3 | 37 | 114 | 60 | 96 | 0.1261 | 0.1815 | 1.5997 | 30 | 0.5343 | 0.0777 | 0.1454 |
| Nadh4L | 38 | 246 | 156 | 289 | 0.1509 | 0.2479 | 2.3937 | 36 | 0.574 | 0.1303 | 0.227 |
| Nadh4 | 38 | 513 | 327 | 612 | 0.1517 | 0.2574 | 2.6205 | 38 | 0.6158 | 0.1336 | 0.217 |
| Nadh5 | 38 | 1440 | 929 | 1880 | 0.1535 | 0.2756 | 3.0058 | 38 | 0.612 | 0.1534 | 0.2507 |
| Nadh6 | 38 | 384 | 258 | 500 | 0.1599 | 0.279 | 2.7935 | 38 | 0.5942 | 0.1588 | 0.2672 |
| 12S | 37 | 592 | 219 | 349 | 0.0886 | 0.1261 | 1.5888 | 36 | n.a. | n.a. | n.a. |
| 16S | 37 | 939 | 395 | 661 | 0.1008 | 0.1522 | 1.9284 | 36 | n.a. | n.a. | n.a. |

***Sagitta elegans* genomes**
| gene | nseff | lseff | S | eta | thetaW | Pi | D | K | PiS | PiNS | w |
| --- | --- | --- | --- | --- | --- | --- | --- | --- | --- | --- | --- |
| Cox1 | 12 | 1032 | 35 | 36 | 0.0112 | 0.0065 | -1.9245 | 12 | 0.0214 | 0.0011 | 0.0528 |
| Cox2 | 12 | 642 | 35 | 36 | 0.0181 | 0.0126 | -1.3559 | 12 | 0.0456 | 0.0019 | 0.0415 |
| Cox3 | 12 | 774 | 34 | 35 | 0.0145 | 0.0093 | -1.6202 | 12 | 0.0376 | 0.0003 | 0.0067 |
| Cytb | 12 | 1137 | 39 | 40 | 0.0114 | 0.0073 | -1.6408 | 12 | 0.0263 | 0.0007 | 0.0266 |
| Nadh1 | 12 | 234 | 11 | 11 | 0.0156 | 0.0095 | -1.6339 | 8 | 0.0378 | 0 | 0 |
| Nadh2 | 12 | 807 | 30 | 30 | 0.0123 | 0.0075 | -1.7542 | 12 | 0.0265 | 0.0012 | 0.046 |
| Nadh3 | 12 | 321 | 11 | 11 | 0.0113 | 0.0071 | -1.5474 | 9 | 0.0255 | 0.0018 | 0.0701 |
| Nadh4L | 12 | 246 | 8 | 8 | 0.0108 | 0.0054 | -1.9834 | 8 | 0.0163 | 0.0016 | 0.0984 |
| Nadh4 | 12 | 792 | 39 | 43 | 0.0163 | 0.0111 | -1.454 | 12 | 0.0384 | 0.0015 | 0.0385 |
| Nadh5 | 12 | 1533 | 63 | 66 | 0.0136 | 0.0086 | -1.7137 | 12 | 0.0265 | 0.0023 | 0.0857 |
| Nadh6 | 11 | 384 | 34 | 34 | 0.0302 | 0.0199 | -1.5944 | 11 | 0.0638 | 0.0044 | 0.0684 |
| 12S | 11 | 557 | 18 | 19 | 0.011 | 0.0074 | -1.4770 | 11 | n.a. | n.a. | n.a. |
| 16S | 12 | 660 | 9 | 9 | 0.0045 | 0.0025 | -1.8304 | 7 | n.a. | n.a. | n.a. |

***Spadella cephaloptera* genomes**
| **gene** | **nseff** | **lseff** | **S** | **eta** | **thetaW** | **Pi** | **D** | **K** | **PiS** | **PiNS** | **w** |
| --- | --- | --- | --- | --- | --- | --- | --- | --- | --- | --- | --- |
| Cox1 | 5 | 1518 | 607 | 810 | 0.1919 | 0.2264 | 1.3708 | 5 | 0.5928 | 0.0525 | 0.0885 |
| Cox2 | 5 | 642 | 309 | 430 | 0.231 | 0.2805 | 1.6362 | 5 | 0.5827 | 0.1208 | 0.2073 |
| Cox3 | 5 | 774 | 369 | 504 | 0.2288 | 0.2713 | 1.4182 | 5 | 0.5743 | 0.113 | 0.1967 |
| Cytb | 5 | 1137 | 452 | 594 | 0.1908 | 0.2214 | 1.2237 | 5 | 0.5828 | 0.0554 | 0.095 |
| Nadh1 | 5 | 870 | 429 | 577 | 0.2367 | 0.2782 | 1.3389 | 5 | 0.583 | 0.1162 | 0.1993 |
| Nadh2 | 5 | 822 | 497 | 666 | 0.2902 | 0.3418 | 1.3597 | 5 | 0.539 | 0.1957 | 0.3631 |
| Nadh3 | 5 | 315 | 159 | 214 | 0.2423 | 0.2876 | 1.4257 | 5 | 0.6008 | 0.128 | 0.2131 |
| Nadh4l | 5 | 255 | 148 | 202 | 0.2786 | 0.3302 | 1.4111 | 5 | 0.5831 | 0.1588 | 0.2722 |
| Nadh4 | 5 | 1227 | 670 | 911 | 0.2621 | 0.3075 | 1.3243 | 5 | 0.5475 | 0.1579 | 0.2883 |
| Nadh5 | 5 | 1536 | 951 | 1330 | 0.2972 | 0.3591 | 1.5938 | 5 | 0.5617 | 0.2042 | 0.3635 |
| Nadh6 | 5 | 423 | 236 | 343 | 0.2678 | 0.3293 | 1.7528 | 5 | 0.5466 | 0.1792 | 0.3278 |
| 12S | 5 | 583 | 235 | 309 | 0.1935 | 0.2185 | 0.9877 | 5 | n.a. | n.a. | n.a. |
| 16S | 5 | 1126 | 386 | 481 | 0.1645 | 0.1843 | 0.9162 | 5 | n.a. | n.a. | n.a. |

**S4 Table.**
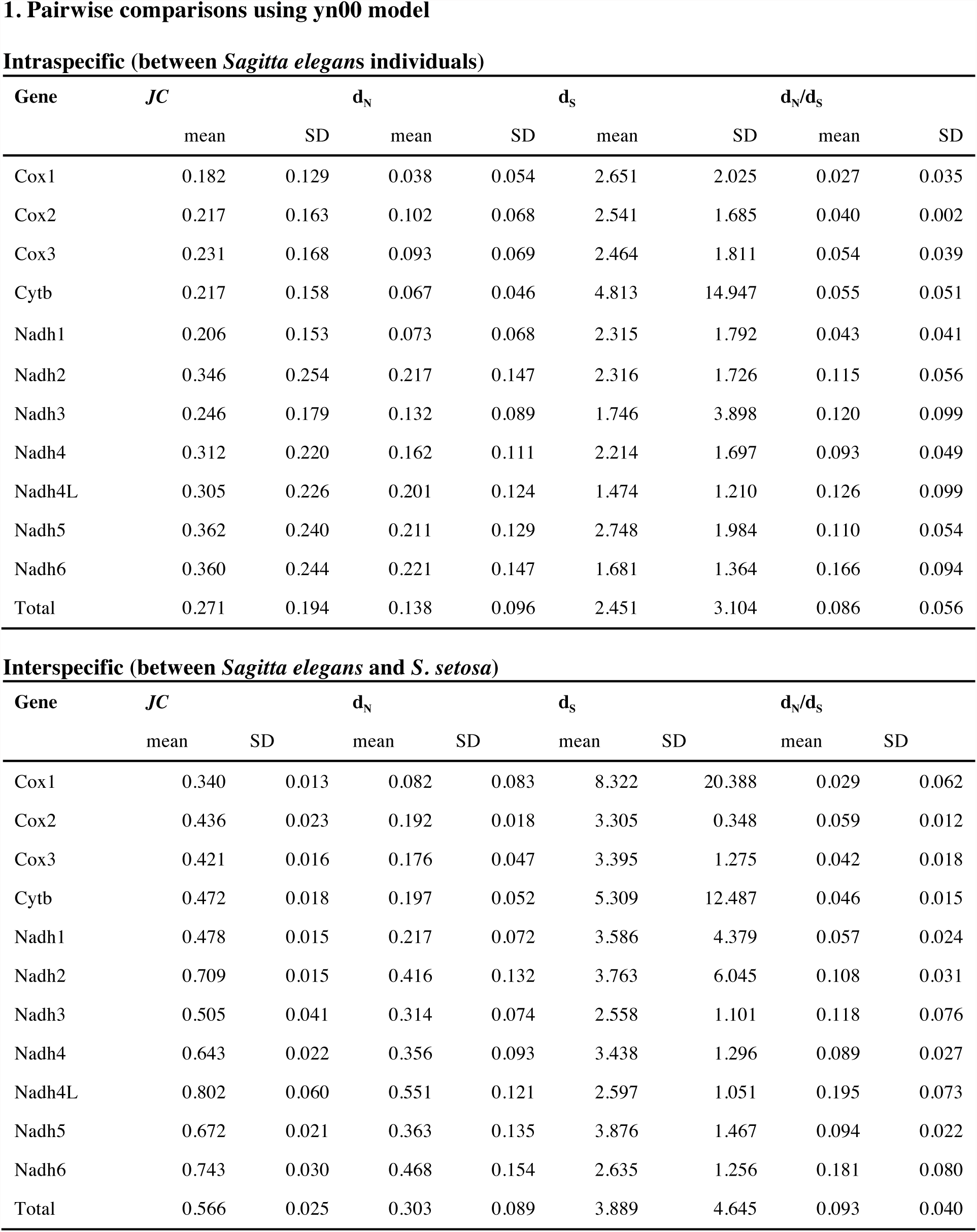
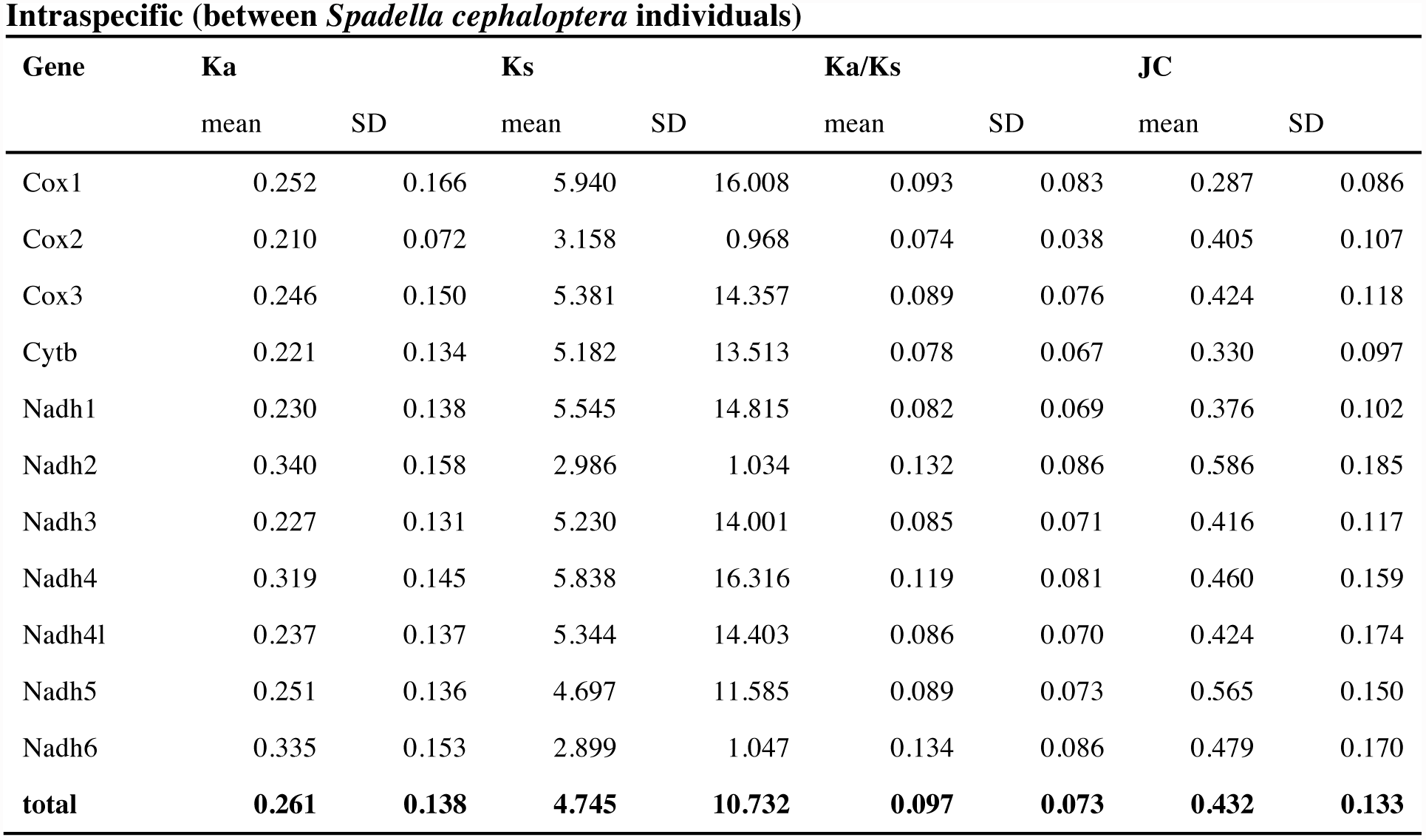
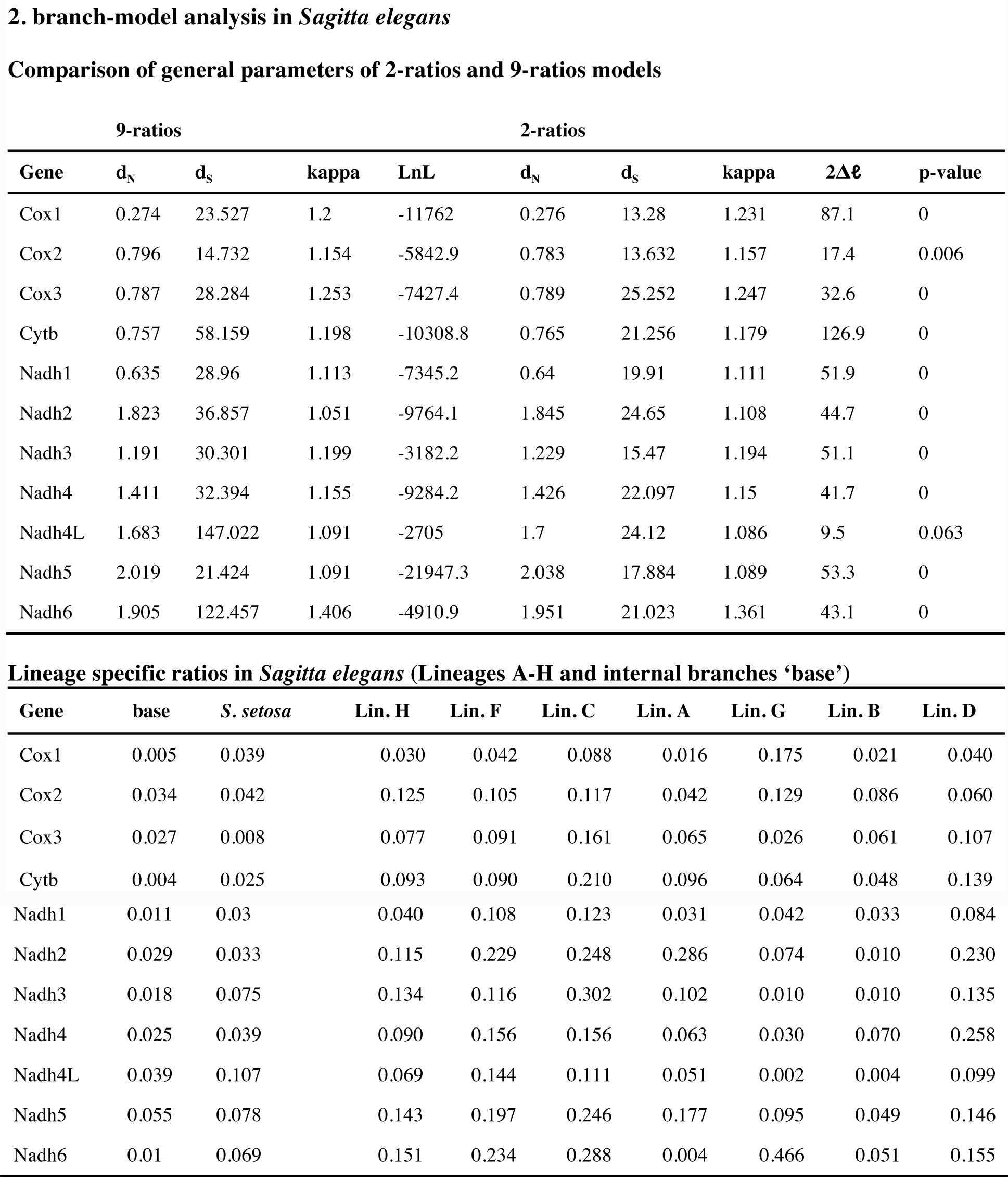
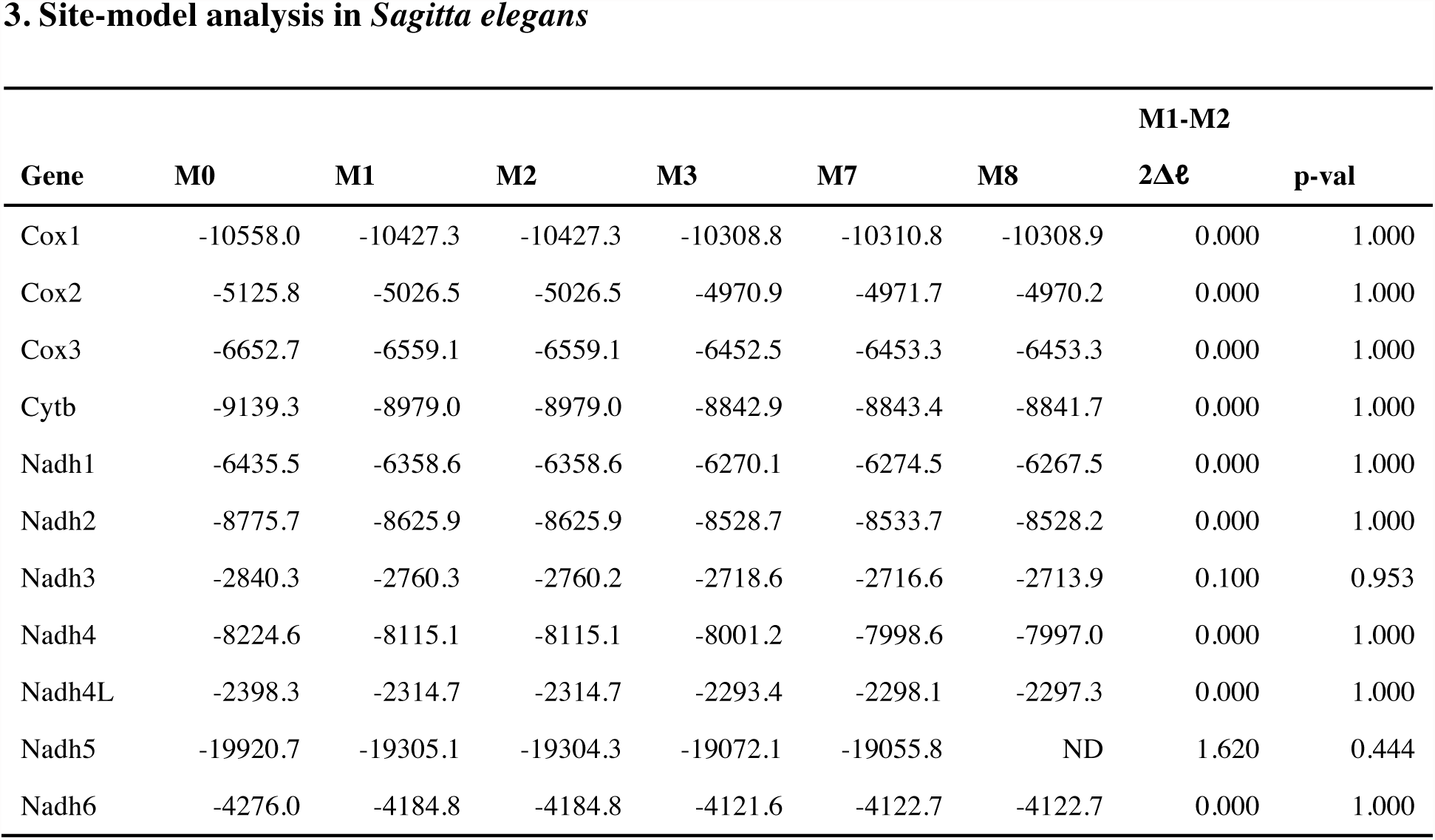
Summary of PAML selection analyses with d_N_, d_S_ and d_N_/d_S_ ratios. (1) pairwise comparisons using yn00 model (2) branch-model analysis with one ratio assigned to each lineage and (3) site-model analysis with results of Likelihood Ratio tests of selection.

**Table S5.**
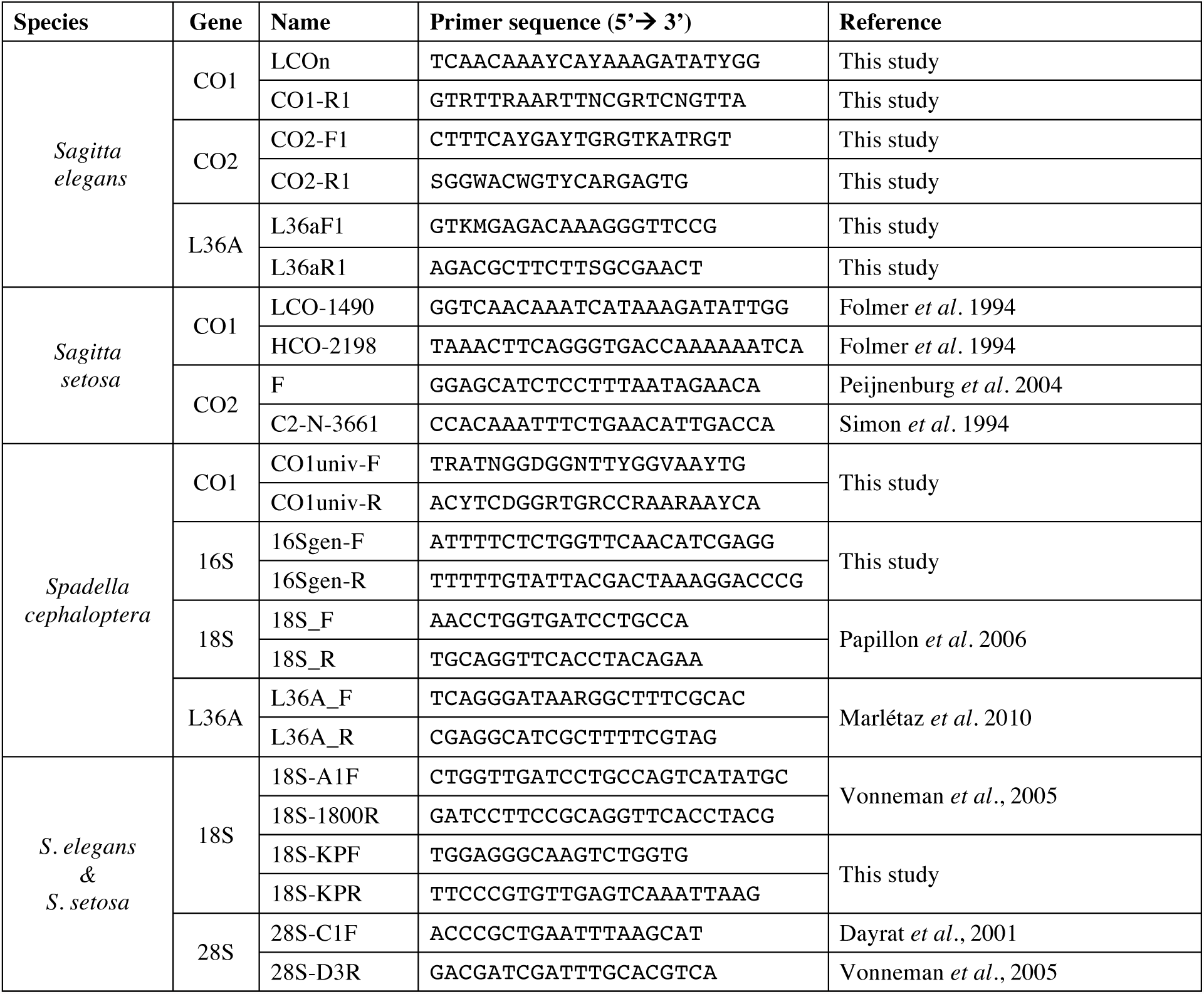
List of oligonucleotide primers used for PCR amplification.

**Dataset S1.** Nucleotide alignments and Maximum-likelihood trees of each genotyped mitochondrial (Cox1, Cox2 and 16S) and nuclear (18S, 28S, L36a) loci in the three investigated species.

**Dataset S2.** Nucleotide, protein alignments and Maximum-likelihood trees of annotated genes in each individual mitochondrial genome.

